# Evolutionary dynamics under phenotypic uncertainty

**DOI:** 10.64898/2026.03.15.711953

**Authors:** Vaibhav Mohanty, Anna Sappington, Eugene I. Shakhnovich, Bonnie Berger

## Abstract

Classical population genetics has largely relied on the same stochastic differential equations (SDEs) for over 60 years to describe evolutionary dynamics. However, these SDEs ignore the fact that phenotype heterogeneity and noise are ubiquitous in biological systems from bacteria to cancers. Here, we develop Probabilistic Phenotype Genetics (ProP Gen) theory as a mathematical framework for evolutionary dynamics under phenotypic uncertainty. Our newly introduced class of SDEs show that, remarkably, phenotypic uncertainty causes many central tenets of classical population genetics to break down, such as invariance of evolutionary dynamics to global shifts in absolute fitness at fixed population size. We revisit the long-studied problem of valley crossing in rugged fitness landscapes, discovering that lower-probability, high-fitness “phenotypic bridges” can substantially accelerate fitness valley crossing even at low mutation rates. We show our theory also explains a paradoxical concept we call “phenotypic buoying” whereby low-fitness phenotypes can exist at surprisingly high frequencies when carried by a high-fitness phenotype which acts as a source. ProP Gen theory uncovers complex phase diagrams of simultaneous coexistence between genotype-phenotype pairs due to phenotypic buoying, which we derive analytically exactly and verify numerically. Notably, the standard Wright-Fisher branching process is unsuitable for probabilistic phenotype numerics because it cannot correctly incorporate phenotypic uncertainty. Thus, we develop a more general, experimentally-inspired discrete-time simulation algorithm, `Probabilistic Serial Dilution (ProSeD)`, which allows for overlapping generations, phenotypic noise, and stochastic phenotype switching. Finally, we show that the new diffusion limit of population genetics recapitulates the experimentally observed resuscitation and partitioning dynamics of bacterial “persister” strains. ProP Gen theory offers promise for describing and predicting the evolutionary dynamics of cancers, which have been empirically observed to exploit phenotypic uncertainty to evade treatment.

## 1 Introduction

Population genetics has provided robust descriptions of the stochastic fate of lineages over time, enabling both forward-time predictions of evolutionary outcomes and backward-time inference of evolutionary histories. The *diffusion limit of population genetics*, a stochastic differential equation (SDE) proposed by Kimura^1^ over 60 years ago, has had a substantial impact in computational biology and genomics, enabling evolutionary simulation methods and laying the groundwork for many central results in genetics^1–8^. The backward-time formulation of this SDE is fundamental to coalescent theory and phylogenetics^9–12^. By explicitly modeling selection-mutation balance and stochasticity via genetic drift, the diffusion limit of population genetics explains how the shape of a fitness landscape impacts the frequency trajectories of genotypes over time. In doing so, this SDE enables avenues for algorithmic prediction of fitness effects, forming the basis of many variant effect and demography estimation methods in statistical genomics^13–18^, and continues to have overarching impacts in many broad subfields of computational biology today.

Although the classical SDE provides a powerful mathematical description of how genotype frequencies change over time on a fitness landscape, it assumes that a genotype uniquely determines its phenotype and that this phenotype has a single, fixed fitness value. This assumption is at odds with empirical observations of real biological systems in which populations of a single genotype can give rise to multiple phenotypic states, often stochastically and with distinct fitness consequences^19–33^. Such phenotypic uncertainty is now recognized as a defining feature of evolution in diverse biological contexts, including bacterial persister cells^20,34–36^ and cancer cell clonal phenotypic heterogeneity^28,29,32,33,37–43^; however, little is understood as to the theoretical underpinnings of phenotypic uncertainty or its impact on evolutionary dynamics. In the setting of phenotypic uncertainty, evolution takes place not on a deterministic fitness landscape^44,45^ but rather on one in which a genotype can map probabilistically to multiple phenotypes^46–48^. Thus, the classical diffusion limit cannot represent these probabilistic mappings, nor the evolutionary consequences of phenotypic uncertainty. Accurately modeling evolution in these systems therefore requires extending population-genetic theory beyond deterministic genotype-phenotype assumptions.

Here, we develop Probabilistic Phenotype Genetics (ProP Gen) theory, a mathematical framework for evolutionary dynamics under phenotypic uncertainty of all kinds. In doing so, we introduce the first SDE that captures the nonlinear interactions of selection, mutation, and phenotypic uncertainty, which broadly encompasses phenotypic plasticity^30,32,33,39,49^, phenotypic noise^24,32,50–53^, and stochastic phenotype switching (SPS)^20,22,25,54^. Our theory predicts striking new dynamical phenomena, such as a novel dependence on absolute fitness, stabilization of low fitness phenotypes by phenotypic “buoys,” and improved fitness valley crossing via phenotypic “bridges.” To validate this theory, we introduce an evolution simulation algorithm, `Probabilistic Serial Dilution (ProSeD)`, which can simulate probabilistic phenotype mapping, unlike the classical Wright-Fisher branching process algorithm. `ProSeD` simulations validate all aforementioned theoretically-predicted phenomena, including a case study in which ProP Gen theory accurately recapitulates experimentally observed bacterial persister cell resuscitation dynamics. ProP Gen theory provides a unified framework from which to study the impact of phenotypic uncertainty on evolutionary dynamics, with important implications for bacterial antibiotic resistance and cancer.

### Related Work

Although Kimura’s SDE^1^ is a mainstay of population genetics literature, mathematical modeling of evolutionary dynamics has not caught up with contemporary experimental interrogations of the role of phenotypic uncertainty in evolution. On the one hand, studies have shown overwhelming evidence that phenotypic uncertainty can drive evolution, from bacterial persister cells^20,34–36^ to cancer cell subclone phenotypic heterogeneity^28,29,32,33,37–43^. Empirically observed phenotypic uncertainty broadly manifests as phenotypic plasticity^30,32,33,39,49^, phenotypic noise^24,32,50–53,55^, and/or SPS^20,22,25,54,56^. However, few proposed models of evolution attempt to capture any form of phenotypic uncertainty. We recently introduced a *static* structural model of probabilistic genotype-phenotype (PrGP) maps^46^. Other studies have subsequently built upon our framework^47,48^, but none of these considers time-dependent evolutionary dynamics. Some simple dynamical models of SPS have added linear generators of phenotypic flux to simple exponential growth^22,34,57^, but these models cannot capture the nonlinear coupling of selection, mutation, and phenotypic uncertainty. Furthermore, to our knowledge, no model has distinguished between phenotypic noise at birth and SPS, which becomes relevant to our discussion of bacterial persister resuscitation dynamics in Section 3.4. An overview of related foundational mathematical models and the evolutionary processes they consider (e.g., selection, mutational flux, SPS, etc.) is provided in Table 1.

**Table 1:** Table of foundational papers introducing population genetics differential equations. This work is the first to incorporate explicit nonlinear coupling of selection, mutation, and phenotypic noise.

| Paper/book | Model | Selection | Genetic drift | Mutational flux | SPS | Mutations AB* | Phenotype noise AB* |
| --- | --- | --- | --- | --- | --- | --- | --- |
| Kimura, 1962<sup>1</sup> | Diffusion limit with drift | ✓ | ✓ |  |  |  |  |
| Crow and Kimura, 1970<sup>58</sup> | Diffusion limit with mutations | ✓ | ✓ | ✓ |  |  |  |
| Eigen, 1971<sup>59</sup>; Eigen et al., 1988<sup>60</sup> | Quasispecies equation, infinite population | ✓ |  |  |  | ✓ |  |
| Thattai and van Oudenaarden, 2004<sup>57</sup>; Acar et al., 2008<sup>22</sup> | SPS with growth | ✓** |  |  | ✓ |  |  |
| **This work** | **All of the above, plus phenotype noise AB** | ✓ | ✓ | ✓ | ✓ | ✓ | ✓ |

\*No assumptions on the weakness of selection/mutation. \*\*unbounded exponential growth, no bottleneck/serial dilution. [SPS = stochastic phenotype switching. AB = at birth.]

ProP Gen theory not only models all of these previously studied evolutionary processes, it also captures phenotypic noise at birth. Derived from a microscopic model, ProP Gen theory explicitly captures the simultaneous coupling of selection, mutation, and phenotypic uncertainty, unlike the classical diffusion limit of population genetics^1^ and current models of selection with SPS^22,34,57^. Ultimately, this unification is a crucial step in understanding real biological evolution in which mutation, selection, and many forms of phenotypic uncertainty all coincide.

## 2. Model: A new time-dependent stochastic description of evolutionary dynamics with phenotypic uncertainty

### 2.1 ProP Gen theory

We now introduce the central theoretical result of this paper: a new SDE that captures selection, genetic drift, spontaneous mutations, SPS, mutations at birth, and phenotypic noise at birth simultaneously. As is standard in modern population genetics^3,61–71^, we assume a haploid asexual population of *N* individuals. Each individual is characterized by a genotype *g* ∈ *G* and phenotype *p* ∈ *P*, with each *p* associated with a Malthusian fitness (i.e., exponential growth rate), *X*^(*p*)^. Traditionally, the assignment of genotype to phenotype occurs deterministically such that individuals of phenotype *p* will produce offspring of phenotype *p* exponentially according to *X*^(*p*)^. In contrast, here we assume the assignment from genotype *g* to phenotype *p* is probabilistic. This converts the standard mountain-like picture of a fitness landscape^44,45^ into a “fuzzier” one; the relationship between genotype, phenotype, and fitness is better visualized as a layering of heatmaps (Figure 1A) where each genotype (horizontal coordinate) can map onto any phenotype (vertical tier) according to some probability vector (color) and replicates according to the associated fitness (height).

**Figure 1:**
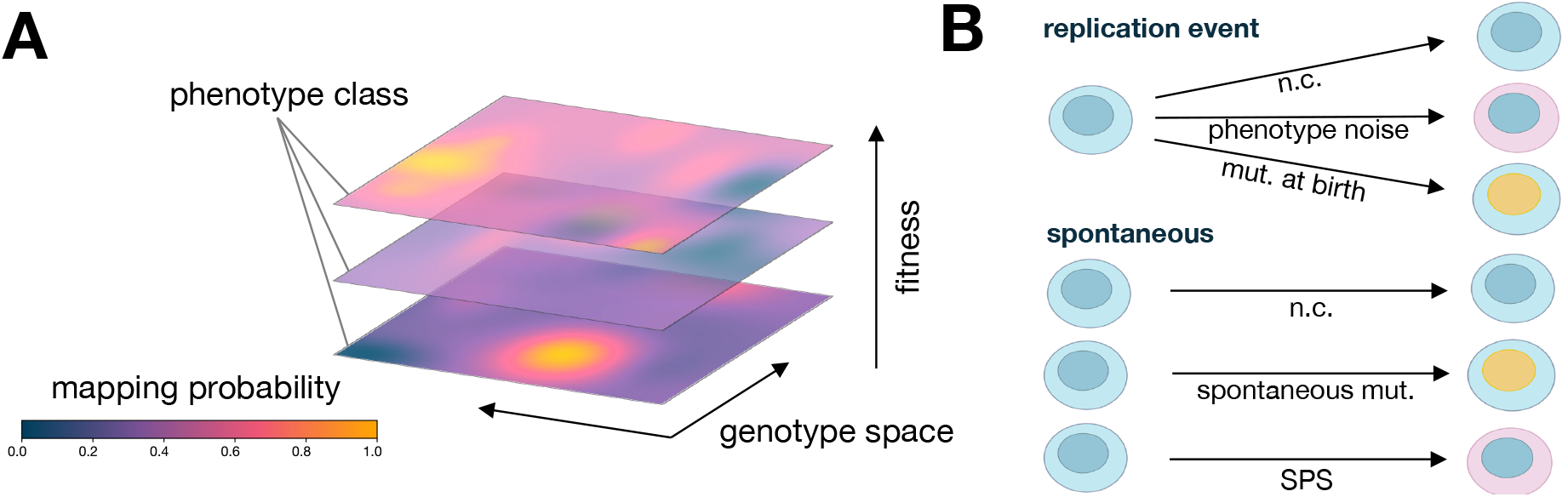
Schematic depictions of probabilistic genotype → phenotype → fitness mappings. (**A**) Diagram of a probabilistic fitness landscape. (**B**) Illustration of types of genotype mutations and phenotypic uncertainty at birth (replication event) and spontaneously. [n.c. = no change; mut. = mutation; SPS = stochastic phenotype switching.]

We next consider a dynamical process through which such haploid populations can evolve over time. At some time *t*, suppose there are 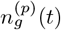 individuals with genotype *g* and phenotype *p*. By the definition of Malthusian fitness, after a short time *δt* we assume all individuals of type (*g, p*) have produced *δtX*^(*p*)^ offspring. Some offspring mutate their genotype *g* to another genotype *h* with probability *µ*_*g*→*h*_, while offspring from all other genotype-phenotype pairs with neighboring genotype *h* can mutate to genotype *g* with probability *µ*_*h*→*g*_. We call this process *mutation at birth* or *mutation during replication*. The mutational probabilities are normalized such that outgoing probabilities sum to one: ∑_*h*∈*G*_ *µ*_*g*→*h*_ = 1.

After offspring genotype assignment during replication, we *noisily* assign a phenotype, possibly with biased inheritance of the parental phenotype, such as in epigenetic inheritance or the partitioning of organelles in eukaryotic cell division. Phenotypic noise is captured by a probability 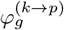, where *k* is the parental phenotype and *p* is the offspring’s possible new phenotype. Thus, during a replication event, both the new genotype and the parental phenotype can influence the outcome of the offspring’s phenotype. We call this process *phenotype noise at birth* and regard it as a distinct type of phenotypic uncertainty. Phenotype probabilities are normalized for every genotype and parental phenotype such that outgoing probabilities of phenotype assignment sum to one: 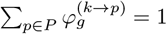

During an organism’s lifetime, we assume heritable mutations can occur, which we call *spontaneous mutations*. The emergence of spontaneous mutations for phenotype *p* takes place at some rate *R*^(*p*)^, with individuals chosen to mutate genotypes from *g* to *h* with probability *m*_*g*→*h*_. Notably, while mutation at replication and spontaneous mutation terms recombine in the deterministic SDE, they are coupled to phenotypic uncertainty in different ways. Phenotypes can also randomly switch during an organism’s lifetime due to SPS, as has been observed baker’s yeast *S. cerevisiae*^22^ and in many bacteria such as *E. coli*^20,36^ and *S. aureus*^25^, which can allow them to adapt to environmental fluctuations. The SPS process takes place at some rate *S*^(*p*)^, with individuals switching phenotypes from *k* to *p* according to their genotype *g* with a probability 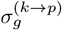. Because phenotypic noise at birth is coupled to both replication rate and mutation rate while SPS is not, the two sources of phenotypic changes lead to differing dynamics.

All of these processes, visualized in Figure 1B, can be combined to write 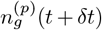 in terms of only abundances 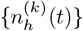 at time *t* as well as the rates and probabilities defined above. We then assume that after time *δt* and all of these simultaneous processes have taken place, the population re-normalizes to the original population size *N*, as if a bacterial culture broth were diluted according to standard laboratory serial dilution protocols. In the limit of small *δt*, small fitness effects, and sufficiently large *N* —the standard assumptions in the classical diffusion limit of population genetics—a *new* Ito SDE emerges for the dynamics of the normalized population fraction 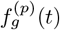:

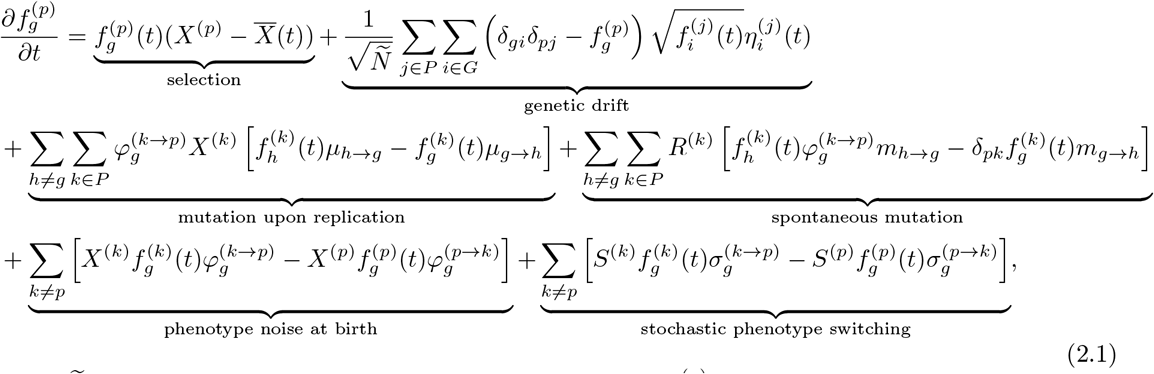

where *Ñ* is population size scaled by generation time units, 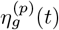 represents white noise with mean 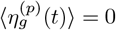 and correlation 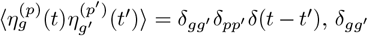 indicates Kronecker delta, and *δ*(·) indicates Dirac delta. We call eq. (2.1) the *Probabilistic Population Genetics (ProP Gen) equation*; it is the central result of this study and its full derivation is included in Appendix A. The ProP Gen equation is the first time a single equation has captured selection, genotype mutations, phenotypic noise, and genetic drift while also distinguishing genetic mutations and phenotype switches at birth from those occurring during an individual’s lifetime. Here, we not only subsume both Eigen’s quasispecies model^59,60^ and Kimura’s diffusion model^1,58^, but we also allow phenotypic noise during birth events to be coupled to *both* mutations and selection. To our knowledge, eq. (2.1) is the first mathematical model that explicitly captures this nonlinear coupling, which we posit is a consequence of biologically realistic and ubiquitous phenotypic uncertainty.

### 2.2 Algorithm for agent-based discrete-time evolution with phenotypic noise

We additionally introduce `ProSeD` (Section B, Algorithm 1), a discrete-time simulation for evolutionary dynamics under phenotypic uncertainty, to address limitations of the widely-used Wright-Fisher branching process. The classical Wright-Fisher branching process assumes a deterministic genotype-to-phenotype mapping and cannot natively incorporate phenotypic noise or SPS. In the `ProSeD` algorithm, when individuals reproduce their offspring undergo potential genotype mutations and then are assigned phenotypes probabilistically, incorporating phenotypic noise. The population is then downsampled to a fixed size (mimicking serial dilution) and the frequencies of all genotype-phenotype pairs are recorded, allowing for the study of their long-term evolutionary trajectories. `ProSeD` can also simulate SPS and phenotypic plasticity through additional probabilistic phenotype and fitness switching steps, respectively. Alongside ProP Gen theory, `ProSeD` allows for a more realistic and comprehensive study of evolutionary dynamics under various forms of phenotypic uncertainty.

## 3 Results

### 3.1 Low fitness phenotypes are stabilized by phenotypic buoys

#### Phenotypic buoys from a simple model of competition

We consider the simplest possible model of competition between multiple genotypes and phenotypes: the case of two genotypes and two phenotypes, which equates to four possible genotype-phenotype pairings. Although simple, such two-state descriptions are accurate approximations in a vast array of biological systems, from binary representations of genotype mutations to gene switches to protein folding energy minima. We will show that the two-genotype, two-phenotype system yields *exact*, analytically tractable long-time population distributions that provide insight into how phenotypic uncertainty can allow less fit phenotypes to survive at high frequencies.

To set up the model, we assign indices of 0 and 1 to each genotype and phenotype, assuming genotypes can mutate between each other with some probability *µ* at each replication event and phenotype assignment only depends on the new individual’s genotype and phenotype assignment probability 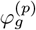. Phenotype *p* has fitness *X*^(*p*)^. This provides a setup as depicted in Figure 2A, with each genotype-phenotype pair depicted as a node in a graph.

**Figure 2:**
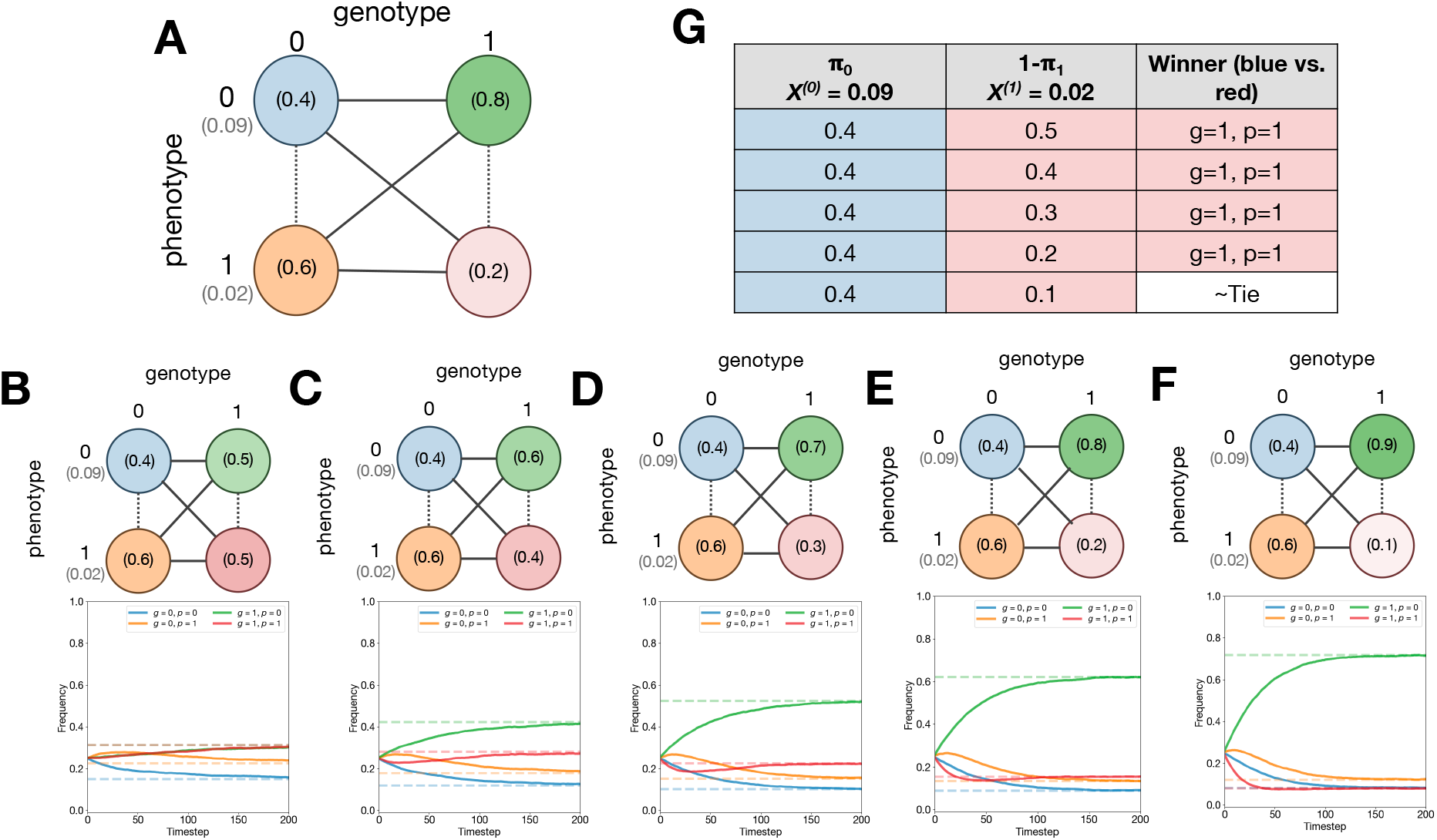
Phenotypic buoys stabilize low fitness, low probability phenotypes. (**A**) Diagram of two genotype, two phenotype model. (**B**-**F**) Diagrams and `ProSeD` simulation results for settings of 1 − *π*_1_ = {0.5, 0.4, 0.3, 0.2, 0.1}. Solid lines is an averaged trajectory from 10 trials *±* SE. Dashed lines are theoretically predicted equilibrium frequencies. (**G**) Table comparing equilibrium frequencies of blue (*g* = 0, *p* = 0) and red (*g* = 1, *p* = 1) nodes.

Working in the infinite population limit (*N* → ∞) with no spontaneous mutations or SPS, we can write down the ProP Gen equations in eq. (2.1) for this system. We then set all time derivatives to zero, since we are interested in the equilibrium population distribution. The mean fitness also becomes time-independent at equilibrium and rearranging the resulting equations, we can write down a 4 × 4 matrix eigenvalue equation for the exact equilibrium frequency vector 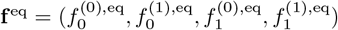. Although 4 × 4 matrix eigenvalue problems are generally difficult to solve, we prove in Appendix C that rank deficiency of the matrix allows for an exact analytical solution for the equilibrium frequencies:

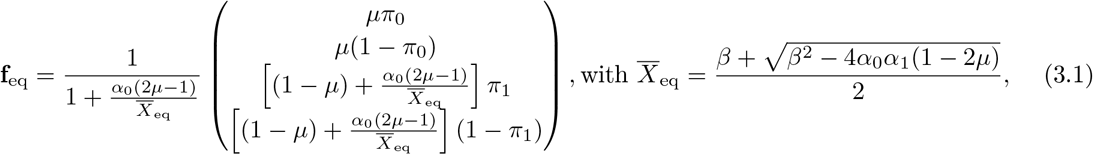

where we introduce the shorthand notation 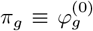 denoting the probability genotype *g* maps onto the higher fitness phenotype 0, 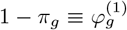 denoting the probability genotype *g* maps onto the lower fitness genotype, 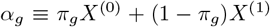, and *β* = (1 − *µ*)(*α*_0_ + *α*_1_). Simplifications of the above vector for special case parameters are discussed in Appendix C.

We then conducted numerical simulations to probe the accuracy and implications of ProP Gen theory. Consider the following thought experiment: can we, for a fixed *π*_0_ = 0.4, intuitively predict whether genotype 1 with phenotype 1 will persist at equilibrium with a frequency higher or lower than genotype 0 with phenotype 0, given some setting of *π*_1_? In Figure 2A, this is equivalent to asking whether the blue node, with mapping probability *π*_0_ and fitness *X*^(0)^, will have a higher equilibrium frequency than the red node, with mapping probability 1 − *π*_1_ and fitness *X*^(1)^.

If the red node has *both* lower mapping probability and lower fitness, naively we might expect that the blue node will exist at equilibrium with higher frequency. Yet, numerical simulations in Figure 2B-F show precisely the opposite result. In these simulations, we ran `ProSeD` (Algorithm 1) with fixed *X*^(0)^, *X*^(1)^, and *π*_0_ while sweeping over *π*_1_. The population was initialized uniformly at random across the genotype-phenotype pairs and simulated until the frequencies reached equilibrium. In each trial, populations equilibrated within 250 dilutions and equilibrium frequencies matched the theoretical predictions from eq. (3.1). Over the entire *π*_1_ probability sweep, summarized in Figure 2G, we found that the red node had higher equilibrium frequency than the blue node in cases where the red node’s mapping probability, 1−*π*_1_, was greater than blue’s, when it was equal to blue’s, *and even* when it was lower than blue’s. Only when 1 − *π*_1_ was reduced to 0.1 did the blue and red equilibrium frequencies approach similar values.

This surprising result is explained by noting that the green node in Figure 2A-F always has high mapping probability *π*_1_ and high fitness *X*^(0)^. Thus, the green node will help absorb total population density from the remaining three nodes (discussed in mathematical depth in Appendix C.3). As it absorbs more population density, it also replicates faster, but during replication phenotype assignment is noisy. As a result, the green node acts as a sink for the blue and orange node’s population density while simultaneously acting as a source for the red node’s population density. At equilibrium, the green node’s frequency ranks highest for *π*_1_ *>* 0.5 and we say that it acts as a *phenotypic buoy* for the red node, which is supported at high equilibrium frequencies despite its mapping probability and fitness being lower than the blue node’s.

This phenotypic buoying phenomenon clearly illustrates why mapping probabilities alone are not trivially predictive of equilibrium frequencies. Mapping probabilities, even when phenotypes are fully non-heritable, are ultimately *conditional* probabilities which represent the mapping probability of a phenotype given its genotype. Mapping probabilities of different phenotypes conditioned on different genotypes require priors on genotype abundances to be predictive.

#### Complex phases of coexistence for genotype-phenotype pairs

We thus see that phenotypic buoys, even in the simplest possible model of competition between two genotypes and two phenotypes, lead to surprising equilibrium frequency distributions. Strikingly, in the presence of genotype mutations, all four genotype-phenotype pairs can persist at equilibrium due to mutations or phenotypic noise. The interplay of these two factors, which are both coupled to replication events, makes predicting the complete ordering of all four genotype-phenotype pairs at long-times a complex problem with 4! = 24 possible permutations.

We next show that the phase diagrams of coexistence between all four genotype-phenotype pairs in any arbitrary parameter regime are exactly tractable, compute the solution analytically, and provide numerical evidence through combinatorially exhaustive parameter sweeps using `ProSeD` (Algorithm 1). The solution follows simply from the equilibrium frequency vector solved in eq. (3.1). Frequency ordering between genotype-phenotype pairs switches at a boundary, which can be calculated by equating any two of the vector elements of **f** ^eq^. There are 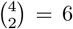 possible equations, which are explicitly written and discussed in Appendix C.

In Figure 3A-C, we plot the theoretical phase boundaries determined from eq. (3.1) as a function of *µ* and *π*_1_. Fixing one of the phase boundary conditions *π*_0_ = 0.5, we make three plots on the (*π*_1_, *µ*) grid, one for each of the conditions *π*_1_ = 0.4 *<* 0.5, *π*_1_ = 0.5, and *π*_1_ = 0.6 *>* 0.5. In Figure 3D-F, we plot the corresponding numerical simulation results for each setting of *π*_0_ where we have empirically determined the equilibrium frequency ordering and colored the space accordingly. For a fixed value of *π*_0_≠ 0, the relative ordering of 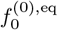 and 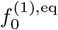 becomes fixed, reducing the maximum number of genotype-phenotype pair permutations by half, resulting in 24*/*2 = 12 possible orderings. The theoretical results show that only 11 sectors are expected for *π*_0_≠ 0, and this is confirmed by the numerical results, aside from noise at boundaries.

**Figure 3:**
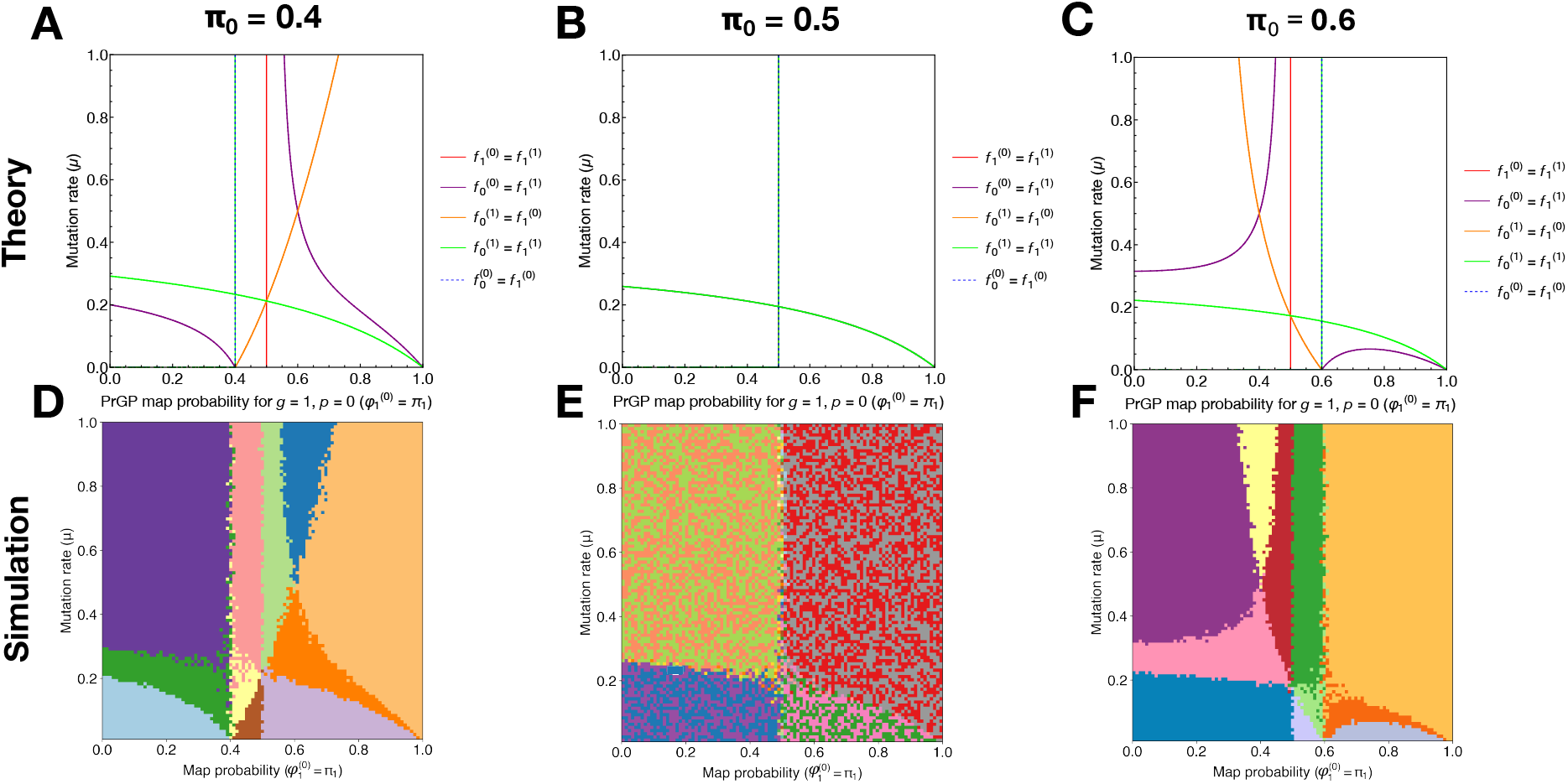
Complex phase diagrams of coexistence between genotype-phenotype pairs at equilibrium. (**A**-**C**) Theoretical phase boundaries for genotype-phenotype pair orderings at equilibrium for *π*_0_ = {0.4, 0.5, 0.6}, from the Prop Gen equation. (**D**-**F**) Corresponding numerical phase diagrams from `ProSeD` for *π*_0_ = {0.4, 0.5, 0.6}.

Altogether, this demonstrates that predicting the ordering of genotype-phenotype pairs at long times is nontrivial. The success of our theory highlights the capability of the ProP Gen equations (eq. (2.1)) to predict the impact of phenotypic noise on evolutionary outcomes. The ProP Gen equations, and phenotypic buoy theory broadly, may shed light on how deleterious mutations can be stabilized in large populations where genetic drift is less impactful but phenotypic noise may instead strongly influence genotype and phenotype abundances. Importantly, although deleterious mutants can be “buoyed” at long times due to mutations, the phase diagrams we have calculated show that standing variation can be generated even in the *absence* of mutations. Moreover, the combined effect of mutations and phenotypic noise may be predictable from our theory.

### 3.2 Phenotypic bridges accelerate fitness valley crossing

Next, we consider a classic problem in population genetics: crossing a fitness valley^3,62,72–75^. Across numerous biological systems, there are cases in which a single mutation is deleterious and thus unlikely to persist, but a second mutation can render the resulting double mutant as fit or even fitter than the wild type. Such double mutations occur in cancers^76,77^, microbes^78,79^, and even drive pandemic waves^80,81^. In the monomorphic regime where mutations tend to fix successively, valley crossing requires extremely rare “stochastic tunneling” in which both mutations simultaneously arise during a single generation. By contrast, in the polymorphic regime populations can spread out across genotypes easily, facilitating crossing. The fitness valley crossing problem has been studied in population genetics over decades^3,62,72–75,82^, but only one study has considered the effect of phenotypic noise^83^ and it does not provide any analytical results which could systematically explain the relationship between crossing time and noise. ProP Gen theory and `ProSeD`, on the other hand, provide a framework for investigating and interpreting valley crossing with phenotypic noise and an *exact* result, despite the complexities arising from multiple strains competing (clonal interference), selection, mutation, and phenotypic noise.

We sought to investigate whether phenotypic uncertainty could allow mutating populations to transiently access neutral or beneficial phenotypes—even ones with low probabilities—thereby accelerating fitness valley crossing. We hypothesized that these *phenotypic bridges* could create alternative routes across rugged landscapes, effectively smoothing or bypassing deep fitness valleys and increasing accessibility of evolutionary transitions that would be improbable under strictly genotype-determined fitness.

Concretely, we consider a system with three genotypes (labeled 0, 1, and 2) and two phenotypes (labeled 0 and 1) with fitnesses *X*^(0)^ and *X*^(1)^ = *γX*^(0)^, respectively, with *γ <* 1. Genotypes 0 and 2 deterministically map the higher fitness phenotype 0, while genotype 1 maps with low probability 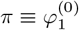 the high fitness “bridge” phenotype and with high probability 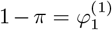 to the lower fitness “valley” phenotype. Mutations occur at birth, making the rate fitness-dependent, with probability of mutation *µ*. In the limit of *π* → 0, the problem becomes the classic deterministic fitness valley crossing problem (Figure 4A) and equilibration across the valley is expected to be slow. For nonzero *π*, phenotypic noise can allow individuals to access the higher fitness phenotype and we investigate whether this increases the speed of population equilibration across the fitness valley (Figure 4B).

**Figure 4:**
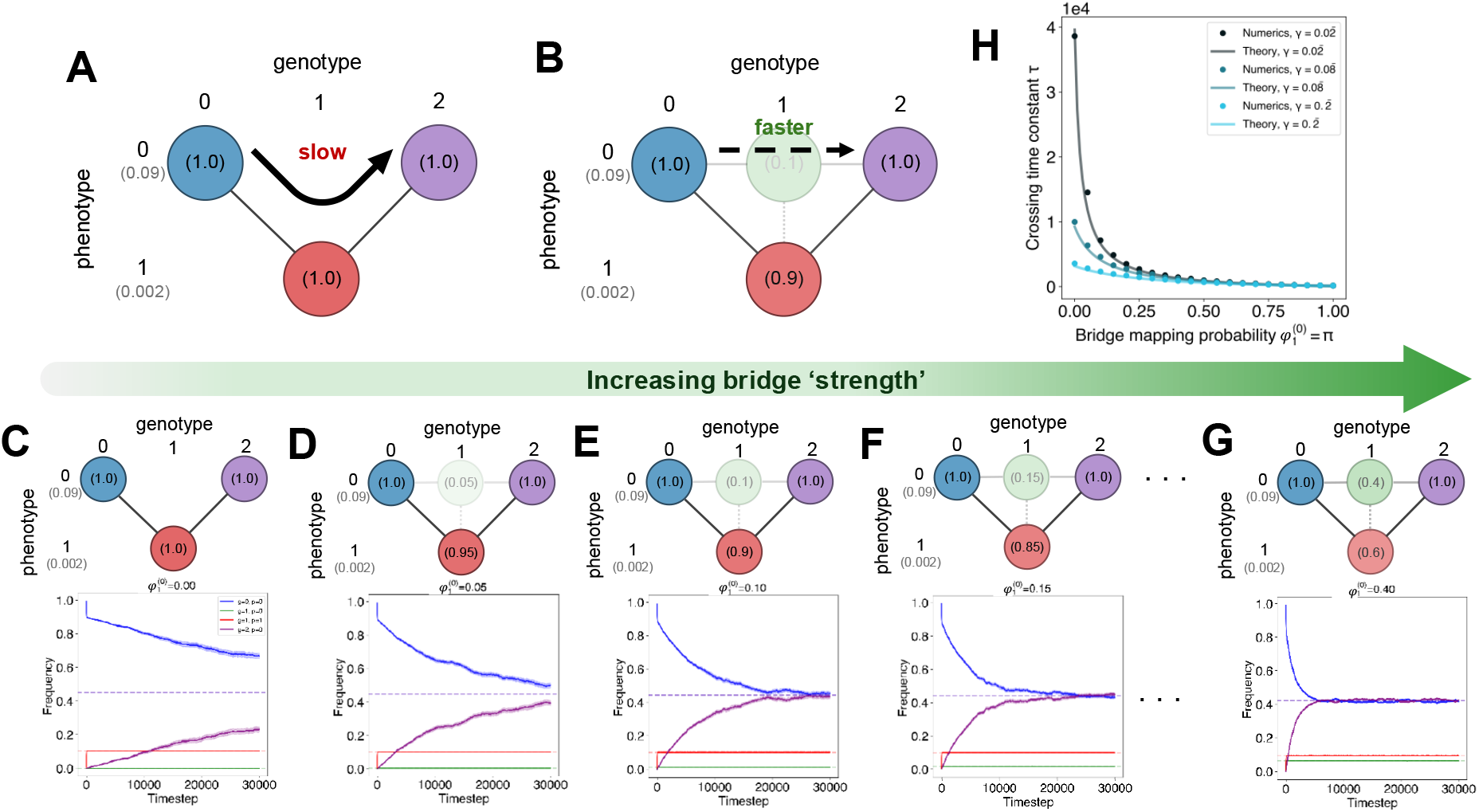
Phenotypic bridges accelerate fitness valley crossing at various depths. (**A**) Schematic of fitness valley crossing in the deterministic genotype-phenotype map setting. (**B**) Schematic of fitness valley crossing in the probabilistic genotype-phenotype map setting. (**C**-**G**) Schematics and time-dependent `ProSeD` simulation dynamics for different bridge strengths *π* = {0.0, 0.05, 0.1, 0.15, 0.4}. Solid lines are the averages of 100 trials *±* SE. (**H**) Bridge crossing/equilibration time constant *τ* versus mapping probability *π* for various valley depths *γ*.

Using eq. (2.1) in the infinite population (*N* → ∞) limit with no spontaneous mutations or SPS, we write the differential equations for each of the four genotype-phenotype pairs with nonzero mapping probability represented in Figure 4B as graph nodes: the starting node *s*, the bridge node *b*, the valley node *v*, and the ending node *e*:

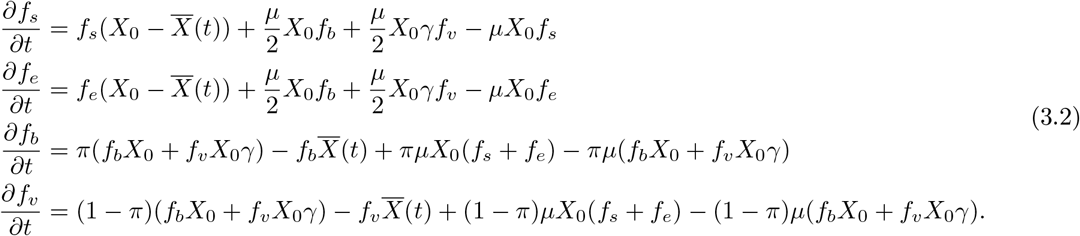

In Appendix D, we calculate the exact equilibrium frequency vector **f** ^eq^ analytically. We then linearize the above nonlinear equations around the equilibrium vector and find the rate of the slowest exponential decay mode by determining the lowest magnitude eigenvalue of the Jacobian. The associated bridge equilibration time constant *τ* is then the reciprocal of the slowest rate; we are able to calculate an exact analytical result:

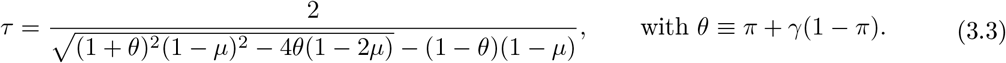

To validate this theory, we run `ProSeD` for various bridge probabilities *π* with an initial population confined to the deterministic node at genotype 0. We then allow the population to relax toward equilibrium; the population fraction on genotype 0 decays and the population fraction on genotype 2 increases towards their respective equilibrium values. Figure 4C-G displays averaged stochastic trajectories from `ProSeD` simulations for various bridge probabilities *π*. We observe faster relaxation toward equilibrium even for very low probability bridges. For instance, even with only a 5% chance of mapping onto the high fitness bridge, exponential decay is notably quicker (Figure 4D). To quantify the relationship between the decay time constant and bridge probability, we use ordinal distance regression to estimate the exponential decay rate for the observed trajectory of *f*_*e*_(*t*), the increasing population fraction on genotype 2. The reciprocal of this rate is the empirical bridge crossing time constant *τ*, which we compare to the theory in eq. (3.3) above.

In Figure 4H, we observe excellent agreement between theory and numerical simulations for *τ* versus *π* across various fitness valley depths *γ*. We emphasize that the theoretical curves are fully determined by the simulation parameters and are not fit to data points, validating our ProP Gen equations (eq. (2.1)) and the derived equilibration time constant *τ* in eq. (3.3) as accurately describing bridge dynamics. Notably, we quantitatively observe a rapid decay of *τ* versus *π* and find that the effect of *π* is larger for deeper fitness valleys (smaller *γ*; Figure 4H).

Importantly, our theoretical predictions and numerical validation demonstrate that even *weak* phenotypic bridges can substantially accelerate fitness valley crossing in the polymorphic regime, where many mutants compete with each other and generations overlap. We are able to show this analytically with highly accurate, zero-fit theoretical curves validated by `ProSeD` simulations. Thus, our theoretical framework provides a robust starting point for modeling how infectious diseases or cancers may exploit phenotypic noise to bypass fitness valleys imposed by immunological or therapeutic defenses.

### 3.3 Mean fitness can decrease with time

#### Absolute, not just relative, fitness impacts population dynamics

Notably, uncertain genotype-phenotype mapping leads population dynamics to depend on absolute, not simply relative, fitnesses. In Appendix E.1, we show mathematically that shifting the entire fitness landscape by some absolute fitness *A* does not transform the selection term in the classic, deterministic SDE but that it does change the probabilistic SDE. Specifically, shifting the fitness landscape by *A* introduces a dependence of the probabilistic SDE selection term on *A*, demonstrating that absolute fitness affects population dynamics under phenotypic uncertainty. Running `ProSeD` evolutionary dynamics simulations at different settings of absolute fitness, while keeping relative fitness constant, demonstrates this behavior. In Figure 5A-C, we see that although shifts in absolute fitness do not affect dynamics in the deterministic setting, they do significantly alter dynamics in the presence of phenotypic uncertainty. Eigen’s quasispecies model has a similar feature, but here we note that phenotypic uncertainty can cause this dependence even in the absence of mutations.

**Figure 5:**
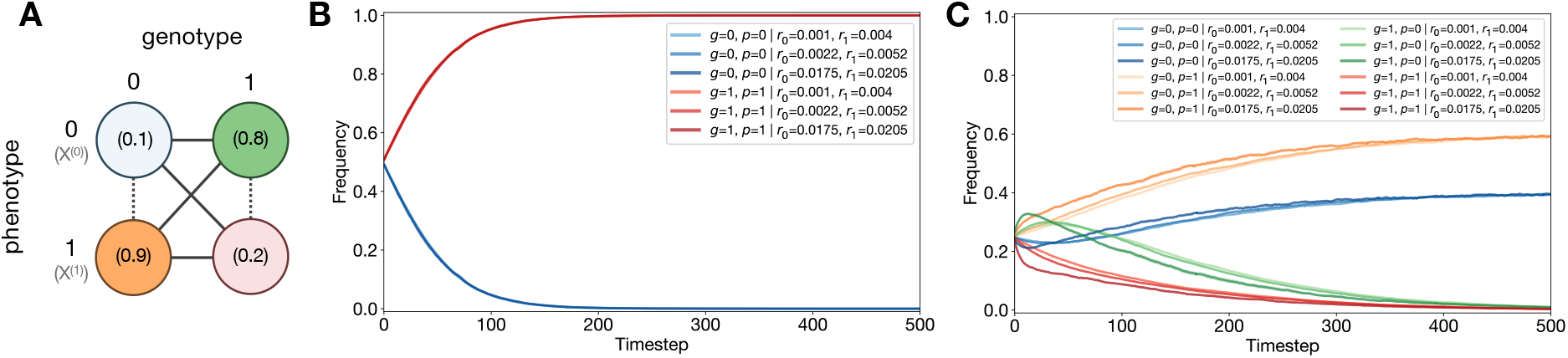
Phenotypic uncertainty leads to dependence on absolute fitness. (**A**) Schematic showing genotype-phenotype pairs and their mapping probabilities, 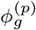, in the setting of phenotypic uncertainty. In the deterministic setting, 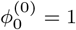 and 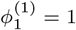. (**B**) Frequencies over time for deterministic genotype-phenotype pairs at different absolute, but the same relative, fitnesses. (**C**) Frequencies over time for probabilistic genotype-phenotype pairs at different absolute, but the same relative, fitnesses. Note *r*_*i*_ ∝ *X*^(*p*)^ as it is the per-generation reproduction probability for a given phenotype *p* used in the `ProSeD` simulations.

#### Phenotypic uncertainty can transiently decrease mean fitness

Moreover, we can construct examples in which the change in mean fitness can be negative over time, even in the setting of no genotype mutations. In other words, phenotypic uncertainty alone can cause mean fitness to decrease. For instance, if we let *X*^(0)^ = 0.09, *X*^(1)^ = 0.02, 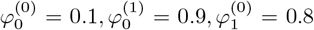, and 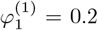 with initial frequencies 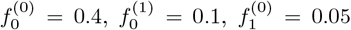, and 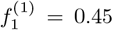, we see in Figure 6A-B and Appendix E.2 that the mean fitness initially decreases. Genotype 0, phenotype 1 (orange) has relatively high mapping probability but low fitness, thus its frequency will be low asymptotically. However initially, its frequency grows due to the relatively high initial frequency of genotype 0, phenotype 0 (blue). The rapid growth of genotype 0, phenotype 1 and non-monotonic behavior contributes to the initial decrease in mean fitness before its eventual increase towards its equilibrium value (Figure 6B). Other evolutionary scenarios (such as specific mutation matrices) can cause transient decrease in mean fitness, but our focus here is to demonstrate that phenotypic uncertainty can cause this decrease even in the absence of mutations.

**Figure 6:**
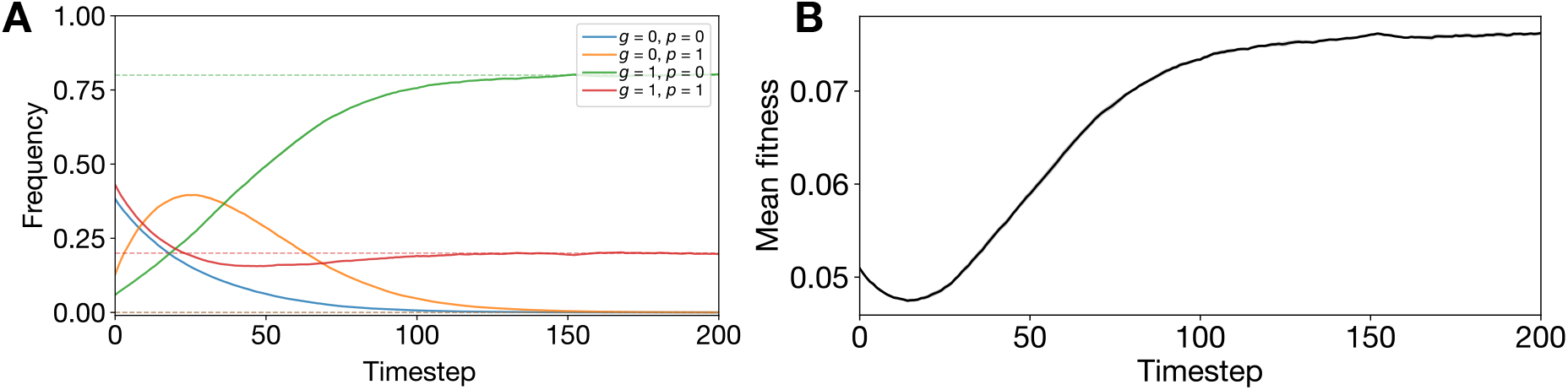
Mean fitness can decrease from phenotypic uncertainty alone. (**A**) Frequency over time of genotype-phenotype pairs with parameter settings detailed in the main text. (**B**) Mean fitness over time for entire population.

### 3.4 Dynamics of bacterial persister resuscitation

We next sought to apply ProP Gen theory to model the resuscitation of bacterial persister cells using experimentally determined SPS parameters^36^, finding that the ProP Gen equations recover empirically observed dynamics^36^. Persister cells are bacterial phenotypic, but not genotypic, variants which are able to survive in the face of environmental stress such as antibiotic treatment. In normal environmental conditions, these cells are less fit than the wild type phenotypic variant and thus reproduce more slowly, but SPS or phenotypic plasticity induced by environmental stress generates a nonzero proportion of these less fit variants. Persister cells enter a state of dormancy and can “resuscitate” after the stressor has been removed.

In Fang and Allison’s^36^ experimental study, dormant *E. coli* and *S. enterica* persister cells resuscitated with an exponential rate after antibiotics were washed away, resuming cell division with varying degrees of success. Damaged persisters carried morphological defects and sometimes were unable to replicate or replicated more slowly and at times “partitioned” to produce damaged offspring and healthy offspring^36^. Successfully reactivated persisters with no morphological damage continued to replicate at baseline rates, producing healthy offspring. The fine-grained time resolution showed that damaged cells were rapidly outnumbered by healthy replicating cells within hours of resuscitation, explaining why previous studies with coarser time measurements had never observed such partitioning.

Here, we show that the ProP Gen equations can be exactly solved for persister resuscitation dynamics, which highlights the distinction between SPS and phenotypic noise at birth. In the experiments, no population bottleneck is imposed, so we ignore genetic drift and remove the mean fitness *X*(*t*) term from the selection portion of eq. (2.1). We write the system of ODEs for absolute abundances {*n*_*i*_(*t*)} of dormant persisters (*P* ), resuscitated cells which fail replication (*F* ), resuscitated cells which are damaged but can replicate (*D*), and resuscitated healthy cells (*H*):

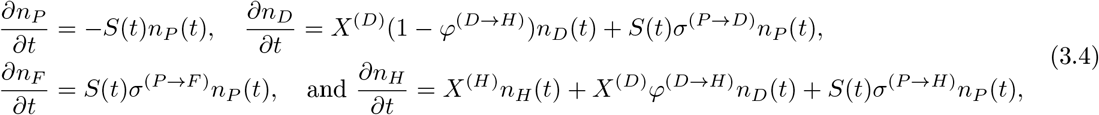

where *S*(*t*) = *αe*^*βt*^ is the exact form of the persister cell resuscitation rate experimentally supported by Fang and Allison^36^, *σ*^*P* →*i*^ for *i* ∈ {*D, F, H*} represents SPS from dormancy to the three resuscitated phenotypes, *X*^(*p*)^ are fitnesses, and *φ*^*D*→*H*^ is the probability that the offspring of a damaged cell is healthy. We show in Appendix F that the above system of equations yields the exact analytical solution

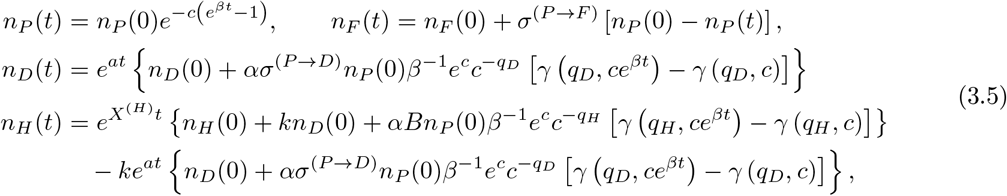

where we have defined shorthand *a* = *X*^(*D*)^ (1 − *φ*^(*D*→*H*)^), 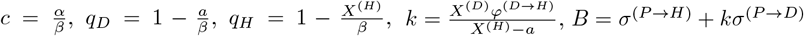, and *γ*(*s, x*) is the lower incomplete gamma function.

In Figure 7, we plot normalized time-dependent trajectories for each of the four phenotypes, provided by eq. (3.5). The dynamics from our theory clearly capture the non-monotonic abundances of failed and damaged persisters, which peak temporarily before disappearing. This transient appearance of abnormal phenotypes accurately recapitulates the experimental findings of Fang and Allison^36^ and explains why prior experiments, which tracked persister resuscitation with coarse temporal resolution, failed to capture these damaged cells. Encouragingly, ProP Gen theory is able to model and predict fine-grained dynamical phenomena observed in experiments stemming from phenotypic uncertainty, showing that the SDE possesses mechanistic fidelity.

**Figure 7:**
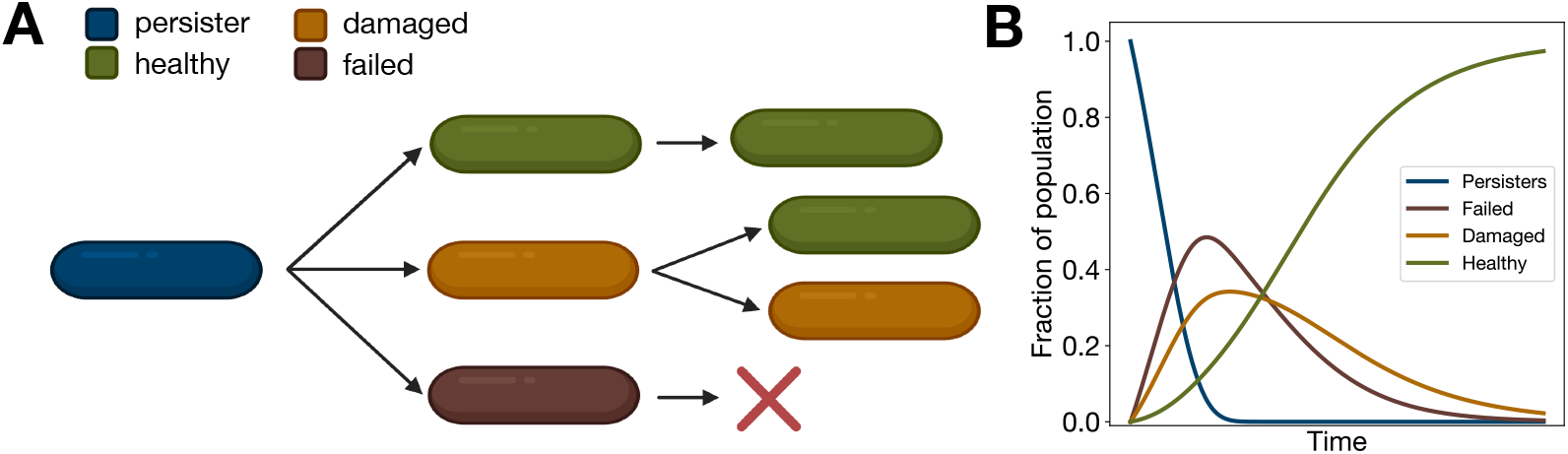
Persister cell SPS case study. (A) Diagram of empirically observed SPS transitions for dormant *E. coli* and *S. enterica* persister cells^36^. All cells have the same genotype. (B) Dynamics of bacterial resuscitation as modeled by eq. (3.5), recapitulating the experimentally observed^36^ transient appearance of damaged and failed resuscitated cells.

## 4 Discussion

We introduced ProP Gen theory, a unified theoretical framework that generalizes the classical diffusion limit of population genetics to explicitly incorporate the interplay of selection, mutation, and phenotypic uncertainty. By allowing each genotype to map to a distribution over phenotypes with potentially distinct fitnesses, switching rates, and replication-coupled noise, our formulation captures a rich set of biological processes that have remained inaccessible to traditional models. Through exact analytical solutions, numerical simulations, and *E. coli* and *S. enterica* persister cell experimental case study, we find that phenotypic uncertainty fundamentally reshapes evolutionary dynamics, altering long-held assumptions about fitness valley crossing and adaptation.

Our theory uncovers two striking new dynamical phenomena caused by phenotypic uncertainty: phenotypic buoying and phenotypic bridging. Phenotypic buoys occur when a high-fitness phenotype stabilizes the persistence of markedly less fit phenotypes, even in the absence of genotype mutations. Exact analytical solutions of the two-genotype, two-phenotype model show that low-fitness phenotypes can dominate at equilibrium when coupled to a strong phenotype source. This finding shows that standing variation in phenotypes can be generated even in the absence of mutations and provides exact results for expected long-term genotype-phenotype distributions when the forces of selection, mutation, phenotypic noise balance each other. In the context of fitness valley crossing, we find that even rare expression of a high fitness phenotype creates phenotypic bridges, alternative transient routes that drastically accelerate equilibration over and crossing between fitness valleys. The exact time constants derived from our diffusion limit match our numerical simulations without parameter fitting, demonstrating the accuracy of our theory. We note the effect of phenotypic bridges is particularly pronounced in deep valleys, suggesting that phenotypic noise may enable highly unlikely evolutionary transitions, including those underlying immune escape in viruses or therapy evasion in cancers. Finally, we find that the experimentally observed transient appearance of damaged cells in bacterial persister resuscitation dynamics can be aptly described by the same ProP Gen model.

Together, these findings highlight that evolution in realistic biological systems cannot be fully understood without explicitly considering phenotypic uncertainty. Populations explore an effective fitness landscape that cannot be fully described by classical deterministic genotype-fitness maps; instead, a PrGP map coupled with ProP Gen theory captures previously unforeseen complexity. We foresee ProP Gen theory offering a promising model by which to explore the effects of phenotypic uncertainty and make testable predictions to be experimentally validated with high-throughput paired genotype-phenotype time series data, which the field currently lacks.

Overall, our work establishes a mathematically rigorous and biologically grounded extension of classical population genetics to incorporate phenotypic uncertainty. By revealing how phenotypic uncertainty reshapes equilibrium and transient adaptive behaviors, ProP Gen opens the door to more complete predictive models of evolutionary dynamics in microbial populations, cancer progression, and beyond.

## Acknowledgments

*Funding acknowledgment*. This work was supported by Hertz Foundation Fellowships (VM; AS), a PD Soros Fellowship (VM), NIH grants R35GM141861 (to BB) and T32GM14427 (to Harvard/MIT MD-PhD Program) from the National Institute of General Medical Sciences. The content is solely the responsibility of the authors and does not necessarily represent the official views of the National Institute of General Medical Sciences, the National Institutes of Health. The authors declare no known conflict of interest.

## Code availability

Code for simulations and plots: https://github.com/vastcollab/ProPGen.

## Supplementary Information for “Evolutionary dynamics under phenotypic uncertainty”

## Appendices

Abbreviations used throughout the appendices include: GP (genotype-phenotype), deterministic genotype-phenotype (DGP) maps, and probabilistic genotype-phenotype (PrGP) maps.

### A Derivation of the PrGP diffusion limit of population genetics as a stochastic differential equation

We derive an Ito stochastic differential equation for dynamics on probabilistic fitness landscapes with an arbitrary number of phenotypes and fitnesses. Here, fitness refers to growth rate given by average replication rate times number of offspring per unit time. Let *G* be the genotype set and let *P* be the phenotype set.

#### A.1 Selection and mutation upon replication, spontaneous mutation, and stochastic phenotype switching

At some time *t* + *δt*, we have

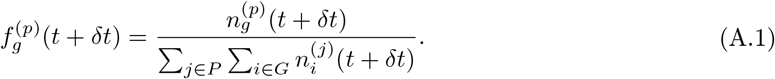

Given fitness *X*^(*p*)^ for a phenotype *p* ∈ *P*, we find that the number of individuals at time *t* + *δt* which have at genotype *g* ∈ *G* which map to phenotype *p* is given below, assuming *δt* is very small:

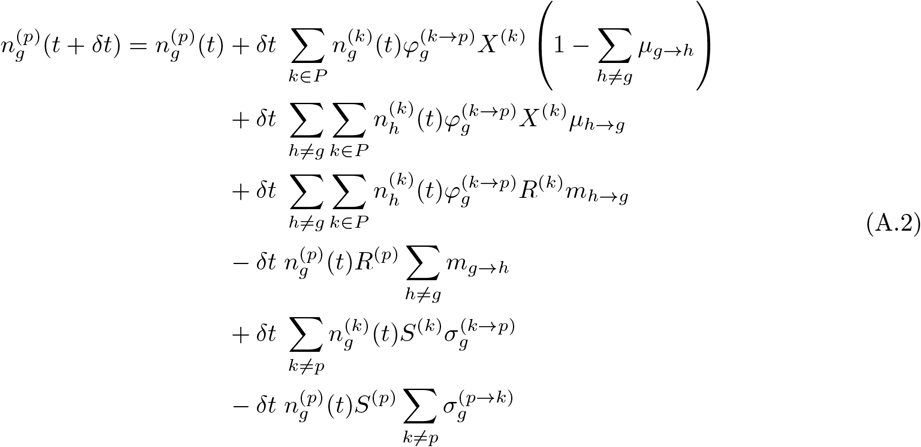

where *µ*_*g*→*h*_ is the probability that a new individual mutates from genotype *g* to genotype *h* at birth, *m*_*g*→*h*_ is the probability that an individual mutates from genotype *g* to genotype *h* spontaneously at a rate *R*^(*p*)^ specified by the starting phenotype *p*, and 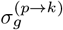 is the probability that an individual changes its phenotype spontaneously from phenotype *p* to phenotype *k* at a rate *S*^(*p*)^ specified by the by the starting phenotype *p* and the genotype *g*. Thus, we have additive contributions from growth of each phenotype, additive and subtractive flux from mutation events which take place at birth (which all depend on 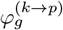 because phenotype assignment occurs at birth but may depend on the phenotype *k* of the parent), additive and subtractive flux from spontaneous mutations which may occur in already existing individuals (which may occur at a rate independent of any growth rates/fitnesses), and additive and subtractive flux from stochastic phenotype switching which may occur in already existing individuals (which also may occur at a rate independent of any growth rates/fitnesses).

We rewrite the equation above, grouping terms into each of these separate contributions

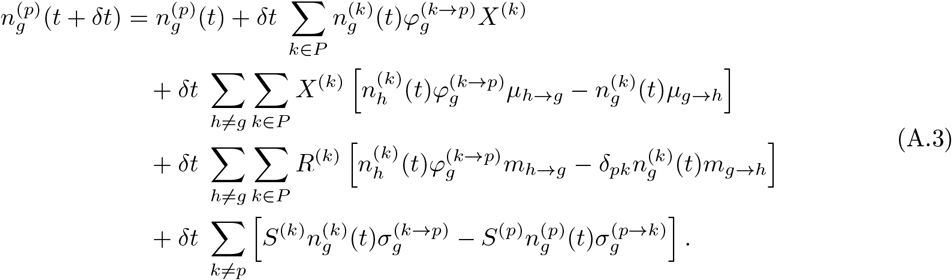

We now compute the denominator of the ratio in eq. (A.1):

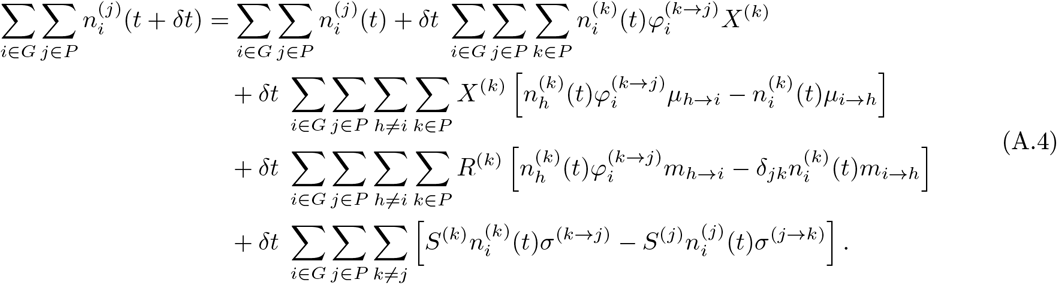

First, we note that the first term normalizes

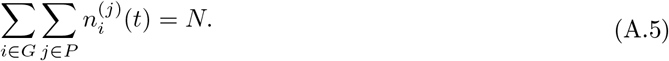

and

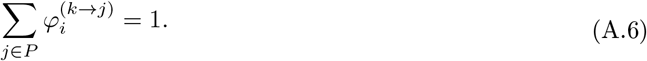

Next, we identify

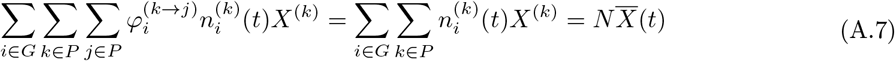

as the mean fitness of the population as a function of time. The third term, which represents mutation upon replication, is simplified

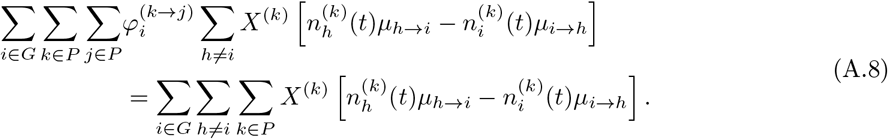

due to the symmetry of the sum

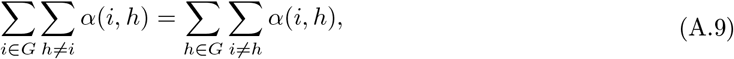

we have that

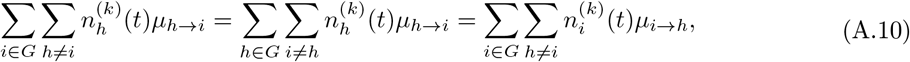

where in the second step we have swapped index labels *h* and *i*. Therefore, the term

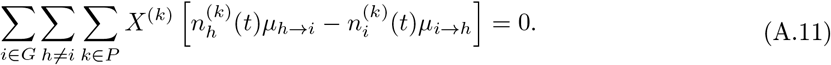

The fourth term, representing spontaneous mutations, is given by

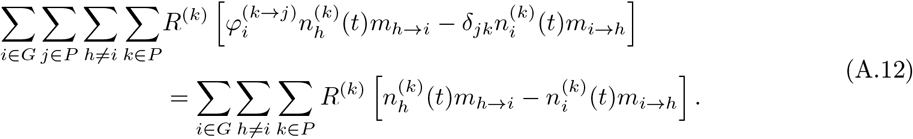

Once again, we have

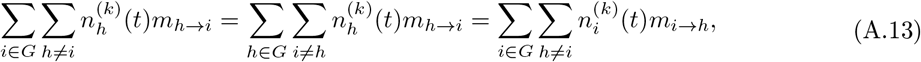

which means

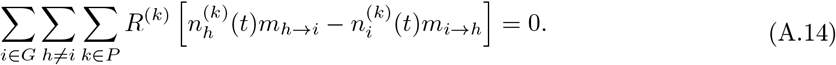

The fifth term, representing stochastic phenotype switching, is

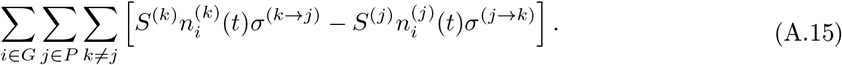

Using the same justification as before,

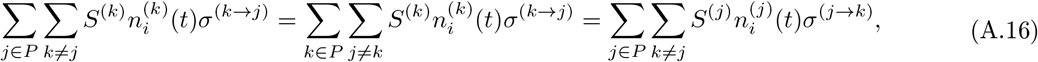

which means

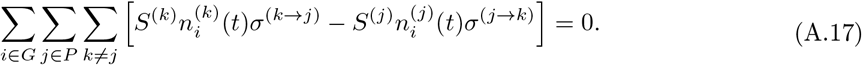

Therefore, the denominator of eq. (A.1) is

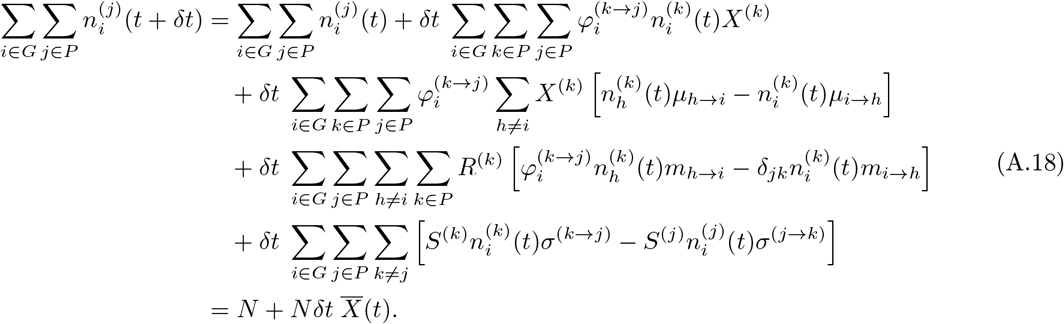

We can now expand eq. (A.1) to 𝒪 (*δt*):

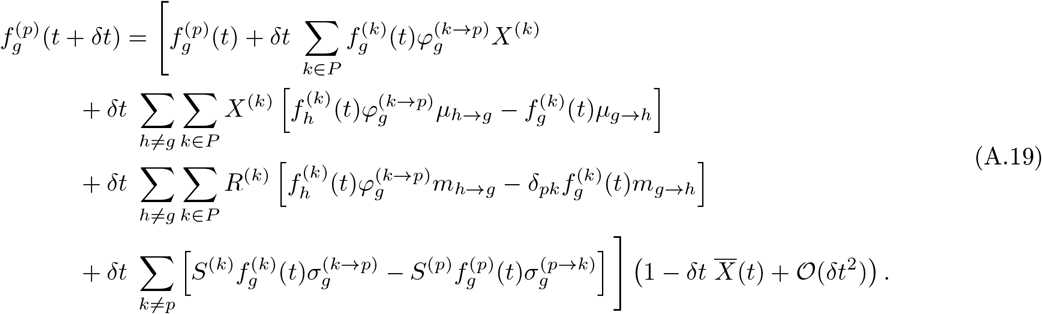

Distributing, we have

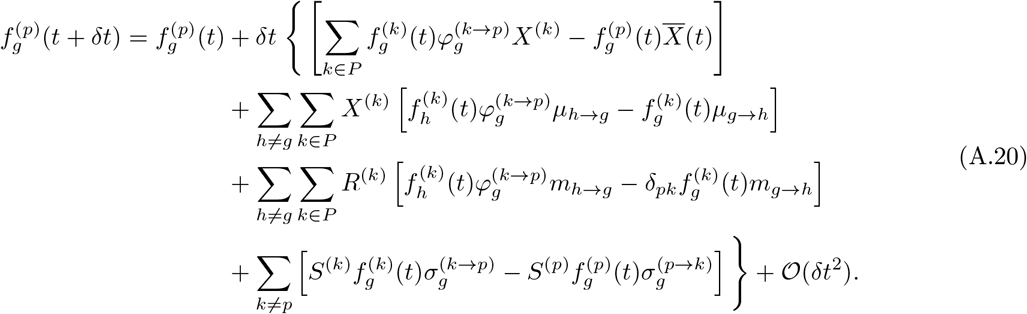

Noting that

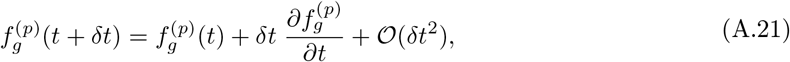

we can write the diffusion limit at infinite population size for dynamics on fitness landscapes with probabilistic genotype-phenotype (PrGP) mapping

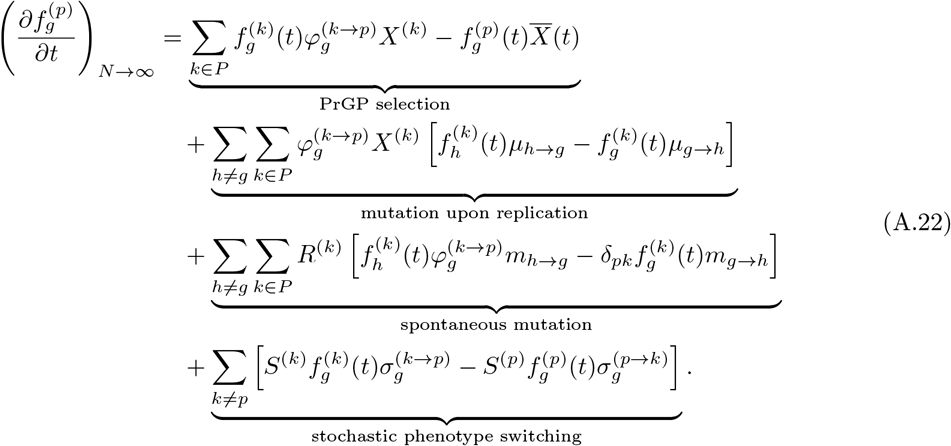

Notably, when 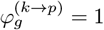 for only one phenotype *p*, meaning that 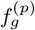 is nonzero for only one phenotype, the exact diffusion limit of population genetics is recovered in the infinite population limit.

Now, we note that we can split the PrGP selection term to two terms resembling classical (DGP) selection (i.e. the replicator term), plus a fitness-dependent phenotype switching term which refers to phenotype noise during replication. To see this, we note

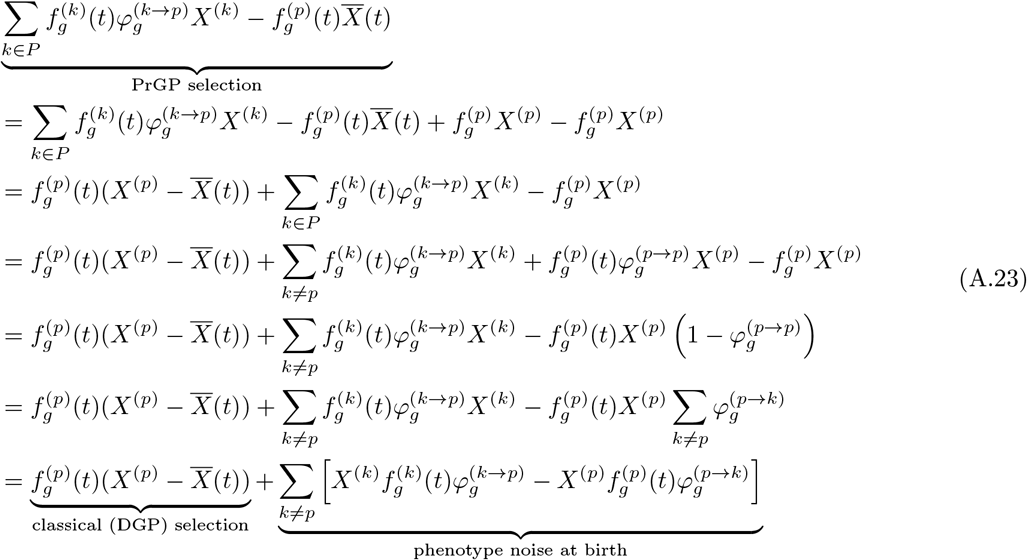

Finally, we can rewrite the diffusion limit at infinite population size as

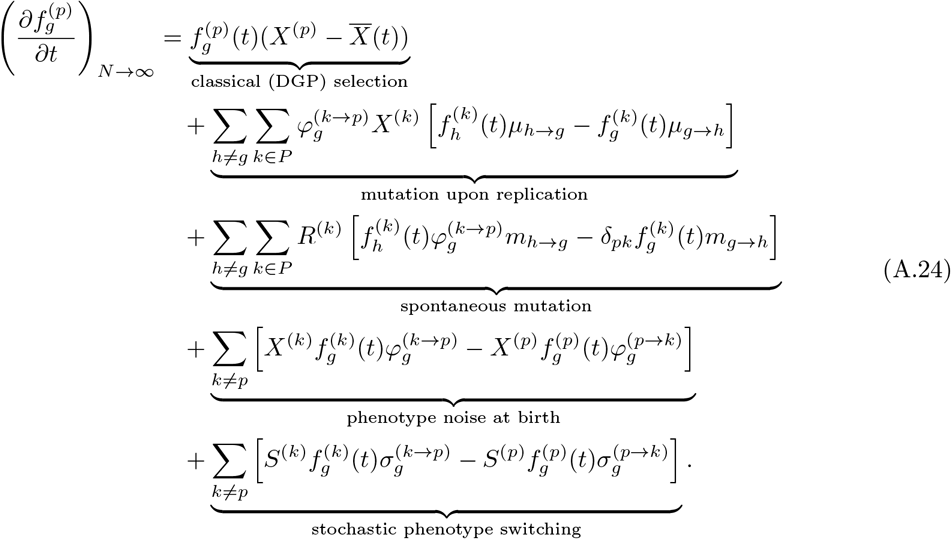

#### A.2 Genetic drift

For finite populations of size *N*, evolutionary dynamics will be subject to genetic drift due to the sampling noise from generation to generation. Generally, in the standard derivation of the (deterministic) Kimura equation, one first assumes that there are no differences in fitness and no mutations at birth. However, as we will discuss in detail in the next session, even if all fitnesses are equal, the absolute fitness determines how many individuals are transferred over from one generation to the next (thus without any need for probabilistic phenotype selection) versus how many individuals are born, all of whom must be probabilistically assigned a phenotype at birth.

We now consider only selection and probabilistic phenotype determination at birth. Thus, the number of individuals with genotype *g* and phenotype *p* at *t* + *δt* will be a random variable sampled from a Poisson distribution

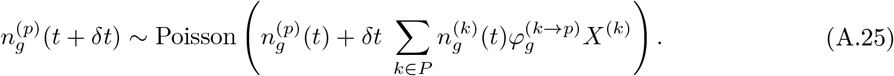

We can write this in the normal distribution approximation

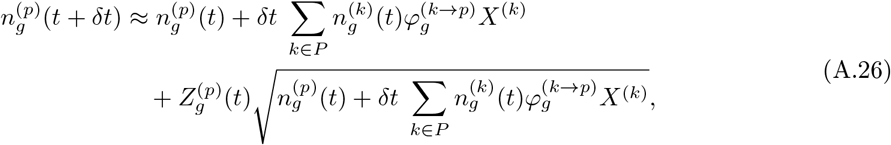

where each 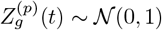 is independent and normally distributed. At some time *t* + *δt*, we have

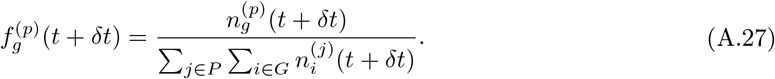

Replacing with the approximation, we have

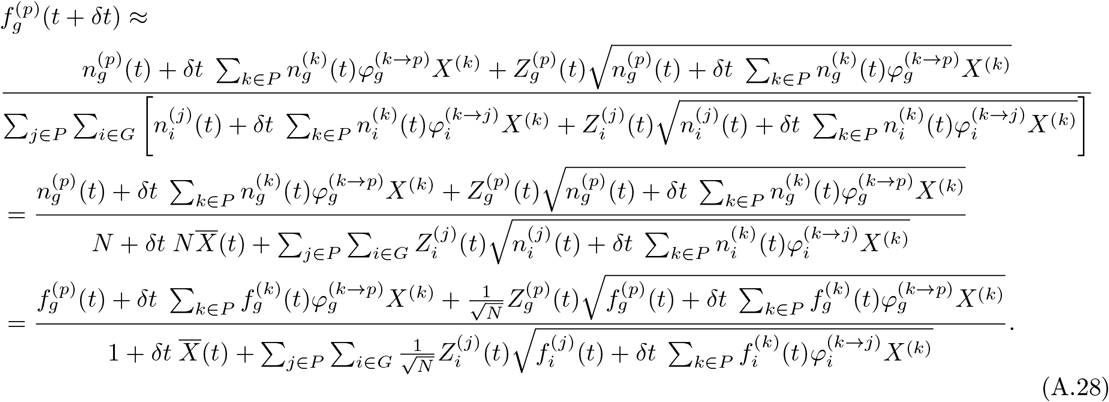

The standard Gaussian 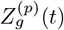 can be related to white noise 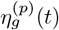 as 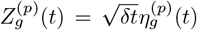 because 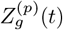 has a variance of 1 over a timestep *δt*. The white noise has mean 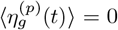 and correlation 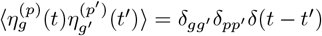. It now follows that

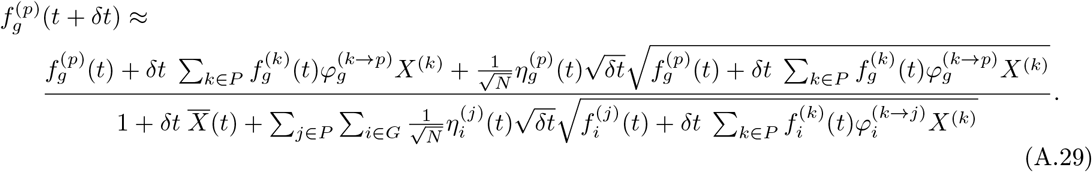

Taylor expanding for small *δt*, we have

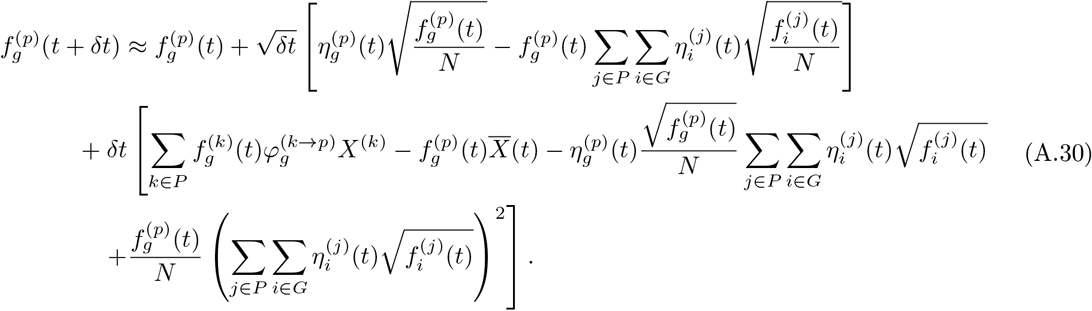

Taking the diffusion limit of small *δt* and large *N*, we ignore the sub-leading order terms which are 𝒪 (1*/N* ), retaining only the terms which are 𝒪 (1) and 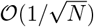 in *N* :

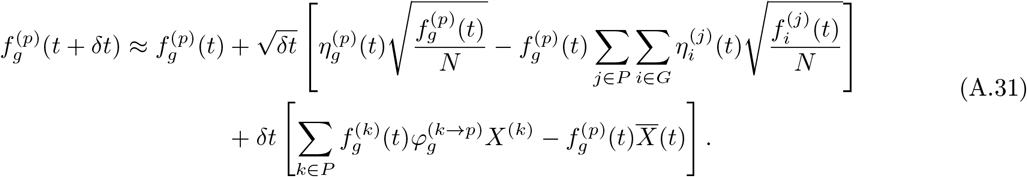

Rearranging, we have

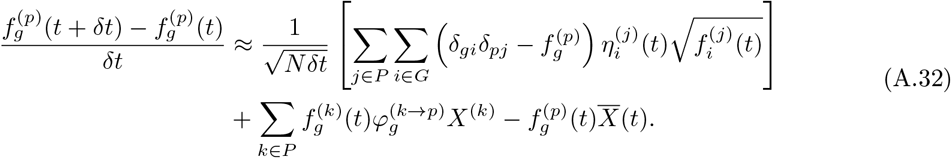

We note that the persistence of 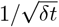 also cancels the units of the white noise, which, from the correlation 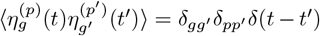 indicates that 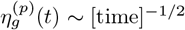. Since we are using *δt* to represent physical time (instead of a non-dimensional time unit such as generations), we let *Ñ* ≡ *Nδt* represent a scaled population size which has been multiplied by the generation time^84^. We now have, in the diffusion limit, the selection term and the genetic drift term:

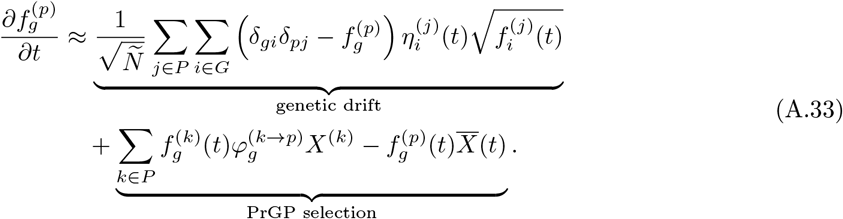

The selection term matches the one derived in eq. (A.24), and the drift term is the same as the diffusion limit of population genetics in the deterministic limit. Thus, we can now write the Itō stochastic differential equation in the Langevin form in the finite population limit, for generation time-scaled population *Ñ* :

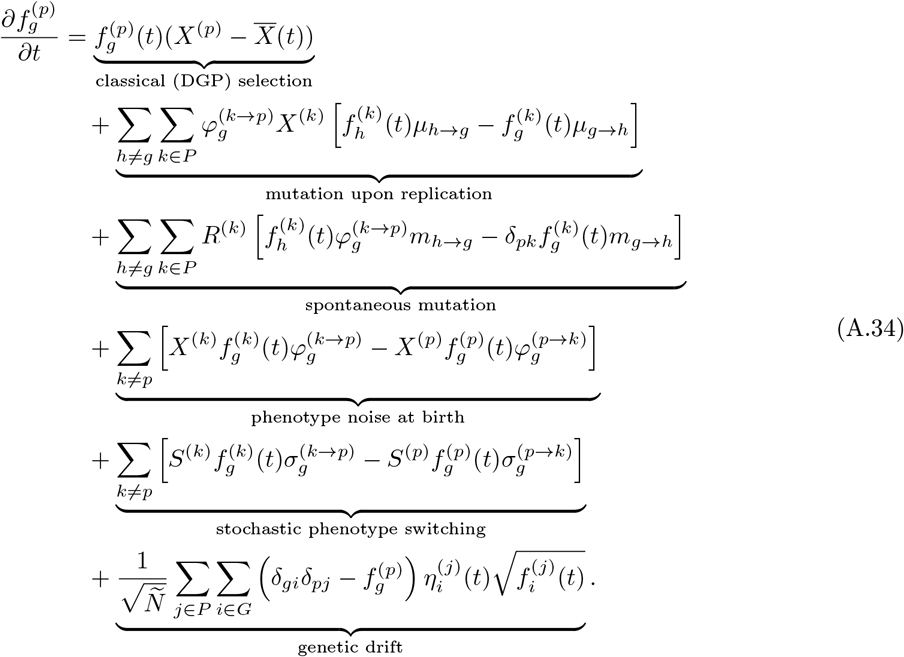

#### A.3 Consistency check: recovering the classical diffusion limit of population genetics in the deterministic genotype-phenotype map regime

We now ask if the deterministic genotype-phenotype (DGP) map—which is the classical diffusion limit of population genetics, equivalent to the Langevin formulation of the multilocus Kimura equation— can be recovered the PrGP diffusion limit which we have just derived. In a standard treatment of population genetics, we would typically only consider selection, mutations, and genetic drift—and maybe recombination, but we are not considering recombination in this treatment. Here, we include both mutations upon replication as well as spontaneous mutations.

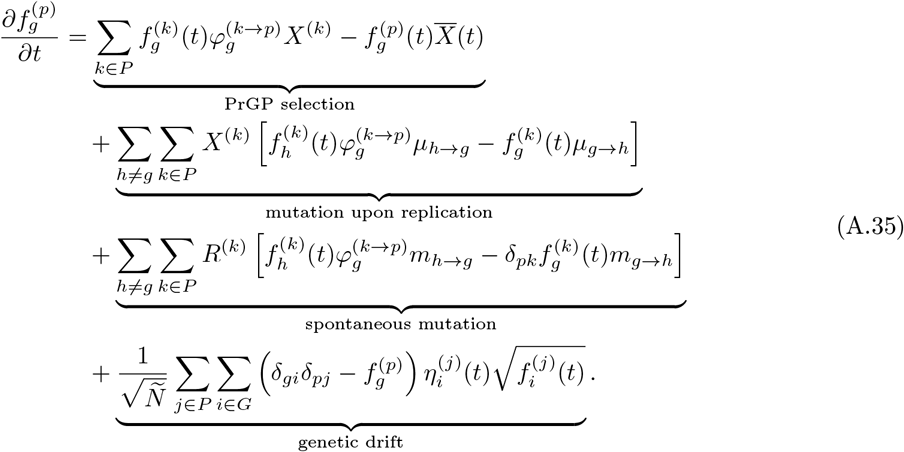

Now, we explore the deterministic genotype-phenotype mapping limit. For convenience, we set phenotype “*g*” to be the phenotype to which genotype *g* maps. This means that each frequency would only have one label: 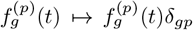, each genotype would have its own fitness *X*^(*p*)^ → *X*^(*p*)^*δ*_*gp*_, and each probability would be a Kronecker delta that is independent of *k* such that 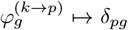. Making these replacements, we have

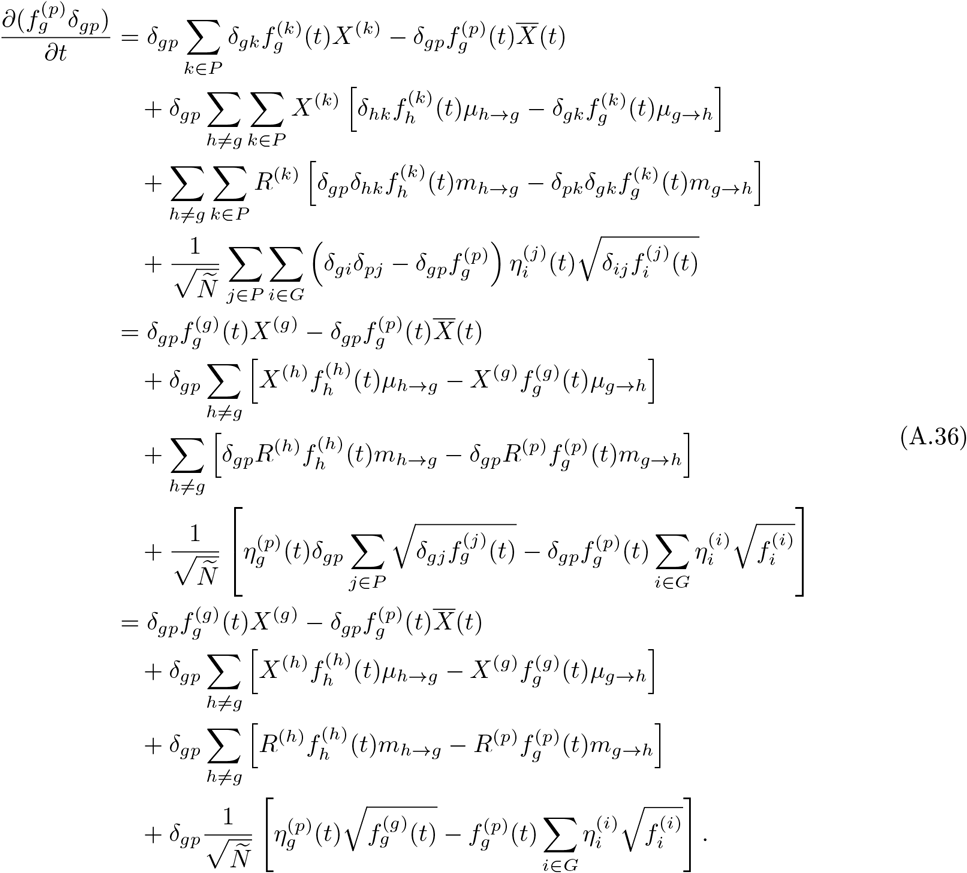

Summing over all phenotypes *p*, and relabeling 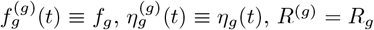, and *X*^(*g*)^ = *X*_g_ we have

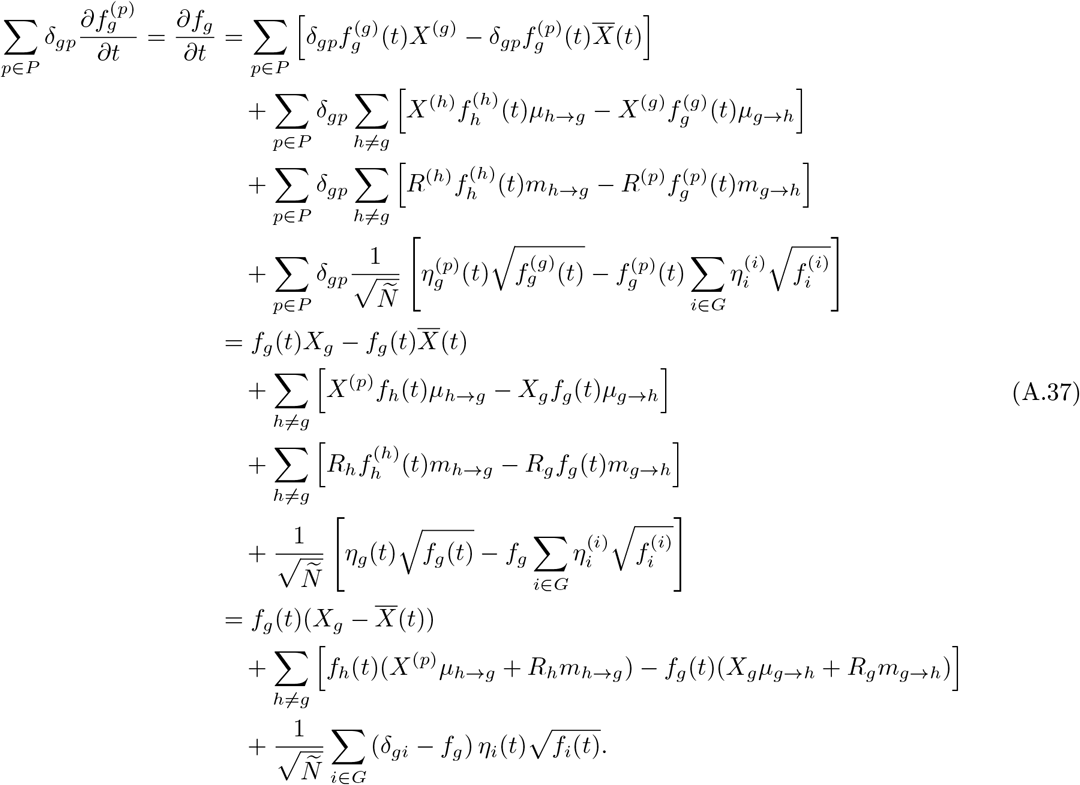

In the final step, we note that *X*^(*p*)^*µ*_*h*→*g*_ + *R*_*h*_*m*_*h*→*g*_ and *X*_*g*_*µ*_*g*→*h*_ + *R*_*g*_*m*_*g*→*h*_ are, respectively, the aggregate incoming and outgoing mutation *rates* which include both mutations upon replication and spontaneous mutations. We have thus arrived at the classical diffusion limit of population genetics, which is equivalent to the multilocus Kimura equation.

### B Probabilistic serial dilution algorithm

Below, we provide pseudocode for the `ProSeD` algorithm, which enables evolutionary dynamics simulations in the presence of phenotype uncertainty.

### C Phenotypic buoys

We now develop the theory of *phenotypic buoys*, where high fitness, low probability phenotypes can support (or “buoy”) the prevalence of low fitness phenotypes to disproportionately high levels at long times.

#### C.1 Calculating equilibrium distributions

We first start with the full PrGP diffusion limit, in eq. (C.1), in the infinite population limit (zero genetic drift), and working in the regime where all mutations happen at birth (and not spontaneously), and there is no stochastic phenotype switching. At long times, the time derivatives of the frequencies will also go to zero, so we have

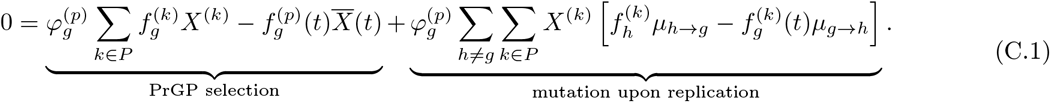

##### Algorithm 1 Probabilistic Serial Dilution (ProSeD)

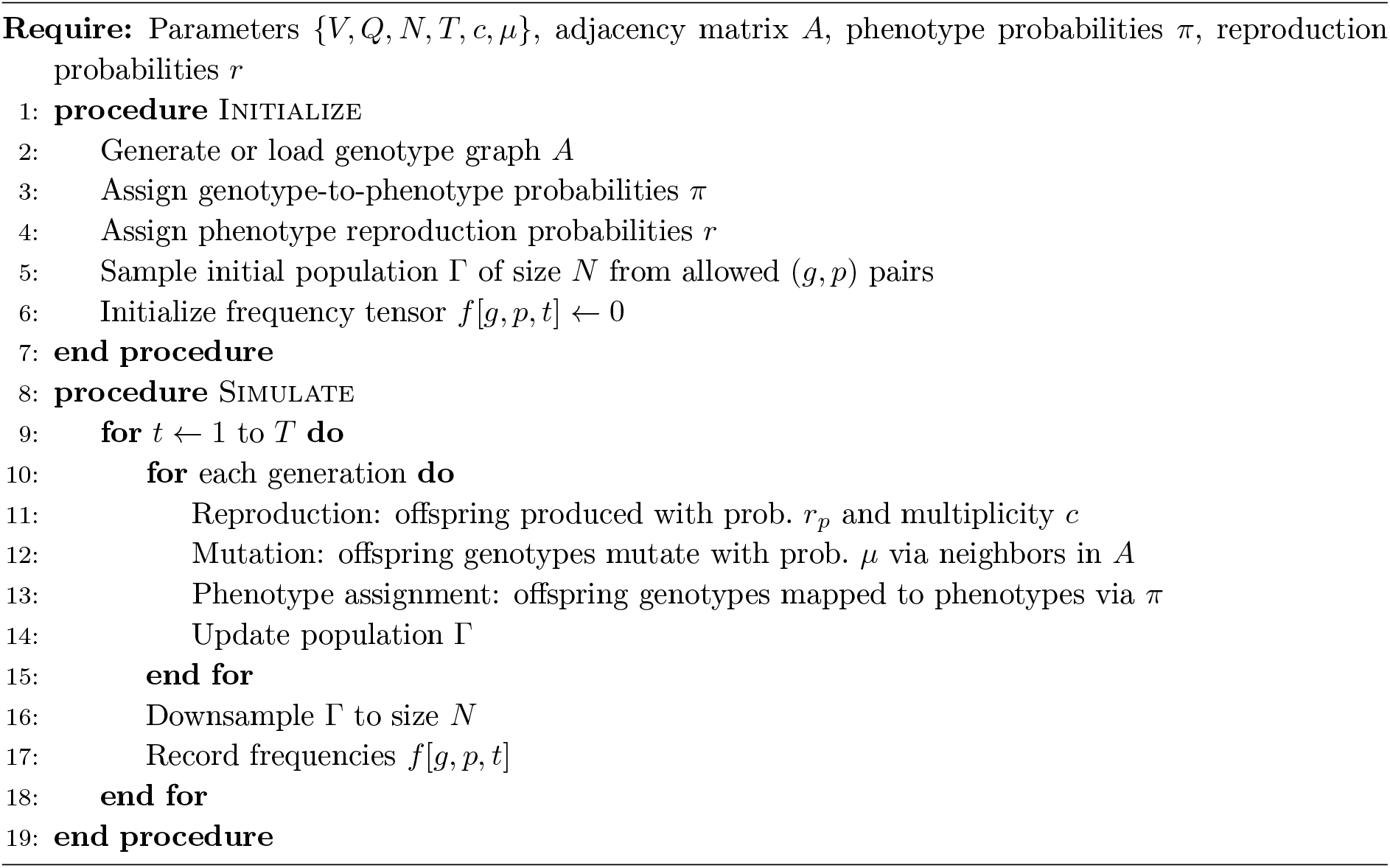

This can be rearranged to

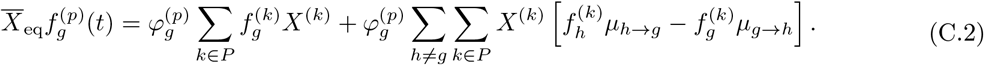

The above equation can be cast as an eigenvalue equation of the form

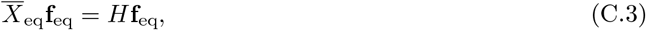

where *H* is a matrix dependent on the fitnesses, PrGP map probabilities, and mutation probabilities. It can be shown using the Perron-Frobenius theorem that there exists a unique equilibrium frequency vector **f**_eq_ with principal eigenvalue equal to the equilibrium mean fitness 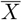. We now use this approach to analytically understand what we describe as a “phenotypic buoy,” where a low-fitness, low-probability GP pair can still persist at equilibrium at unexpectedly high frequencies because it is “buoyed” by a high-fitness phenotype which has the same genotype. To do so, we explicitly construct the matrix form of the above equation for the 2 genotype, 2 phenotype case, which is one of the simplest possible examples that would exhibit this phenomenon.

#### C.2 Exactly analytically tractable example case: phenotypic buoy with 2 genotypes and 2 phenotypes

Suppose we haves two genotypes labeled 0 and 1, and two phenotypes labeled 0 and 1. We can then use the equations above to write the equilibrium matrix relation

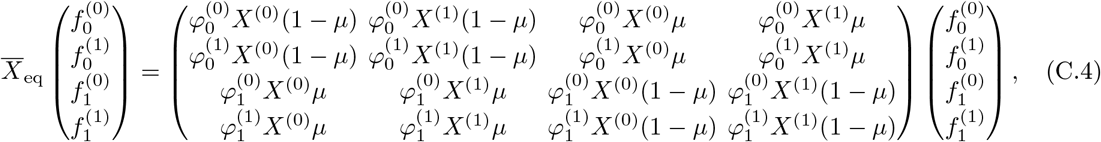

where we have assumed uniform mutation probabilities *µ*. We can write the equilibrium equation in block matrix form

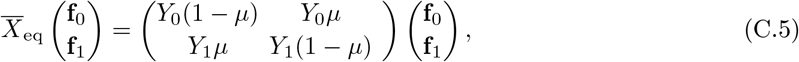

where we defined

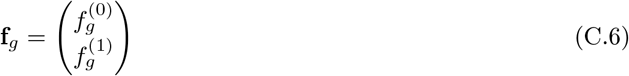

and we can express the matrices as outer product

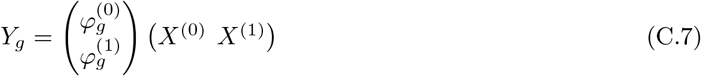

Next, we note that since we only have two phenotypes, we can further denote

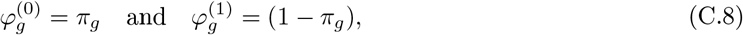

so that we have

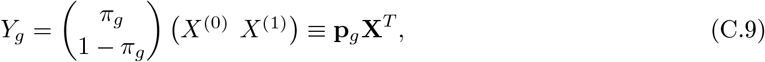

where we have defined vectors

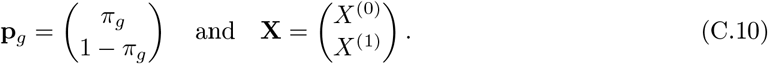

The eigenvalue equation can now be decomposed into two equations:

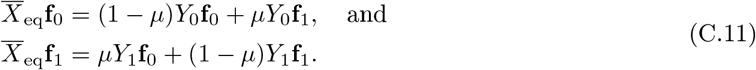

In terms of the outer product decomposition, we can write

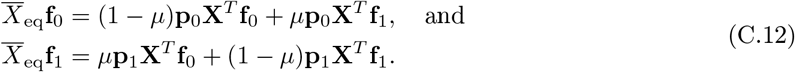

Defining scalar dot products between the fitness and genotype-specific frequency vectors,

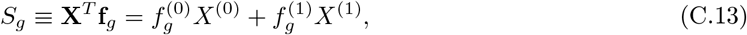

we can write

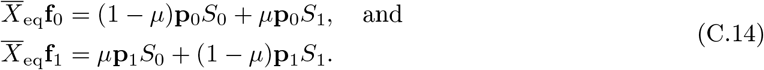

Multiplying both sides by **X**^*T*^ from the left, we have

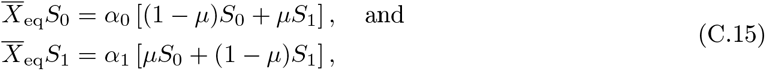

where we have also defined two new scalars which are dot products between the fitness and the genotype-specific GP mapping probability vectors:

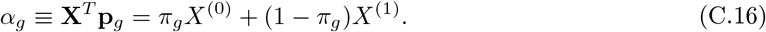

Rearranging the matrix equation, we can now frame the system of equations a new 2×2 matrix eigenvalue problem

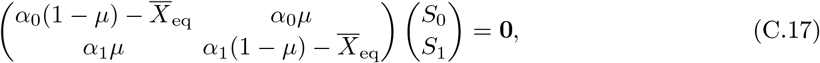

which holds when

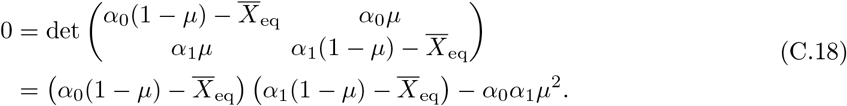

The original 4 × 4 matrix has two eigenvalues which are zero, because the block matrices are each rank-1 (since they are expressible as an outer product). The above quadratic equation provides the two non-zero eigenvalues and corresponding eigenvectors. Solving the quadratic equation, we obtain eigenvalues

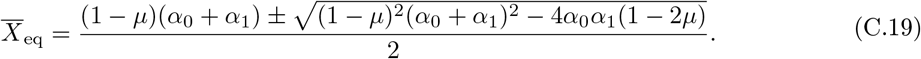

As mentioned previously, the Perron-Frobenius theorem indicates that the principal eigenvalue and its corresponding eigenvector will give us the equilibrium mean fitness and frequency vector, respectively. Thus, we only take the positive solution:

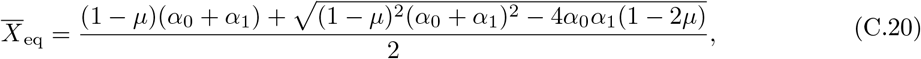

to be the true equilibrium mean fitness. Now, we can use it to find the equilibrium frequency vector. First, assuming mutation probabilities are nonzero (*µ >* 0), we note that

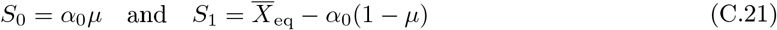

is a valid solution to eq. (C.17). Substituting these values into eq. (C.14), we can obtain unnormalized components of the equilibrium eigenvector

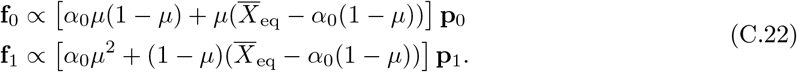

Normalizing and simplifying the frequency vector, we have the complete equilibrium frequency vector expressed as

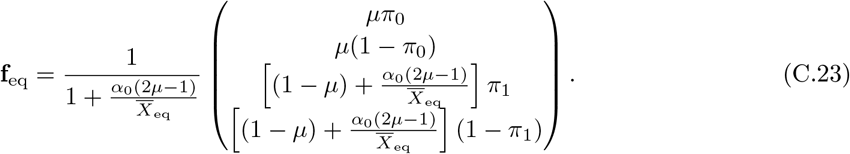

This holds in general when the mutation rate *µ* is nonzero.

#### *Limiting case: zero mutations* (*µ* = 0)

When no mutations are present, the eigenspace becomes degenerate

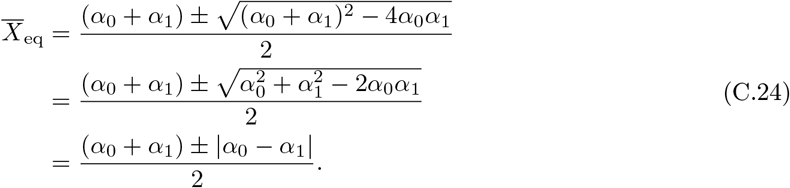

When *α*_0_ *> α*_1_,

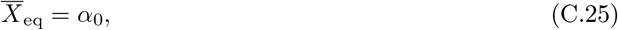

from which it follows that

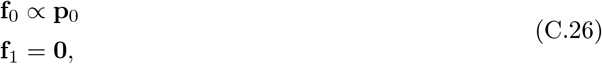

so

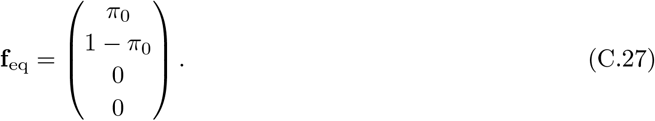

When *α*_1_ *> α*_0_,

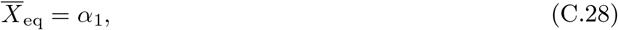

from which it follows that

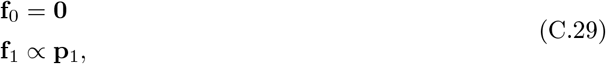

so

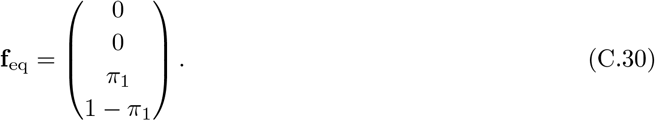

When *α*_0_ = *α*_1_, the principal eigenvector is degenerate and any linear combination of the above two eigenvectors will be a valid equilibrium eigenvector.

#### Limiting case

*zero fitness of phenotype 1* (*X*^(0)^ = 0). We now explore the limiting case where *X*^(1)^ = 0, some simplifications can be made. In this limit,

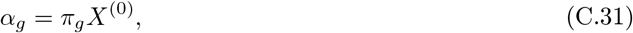

from which it follows that

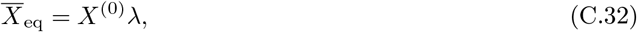

where we define

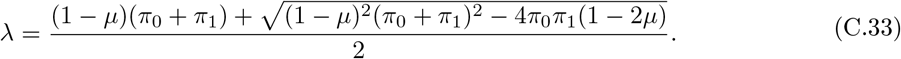

now, we note that the fraction

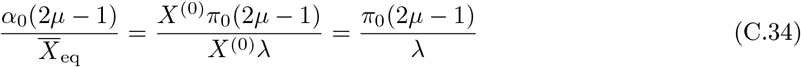

loses dependence on the the fitness of phenotype 0. As a result, the equilibrium frequency vector no longer has any dependence on fitness whatsoever:

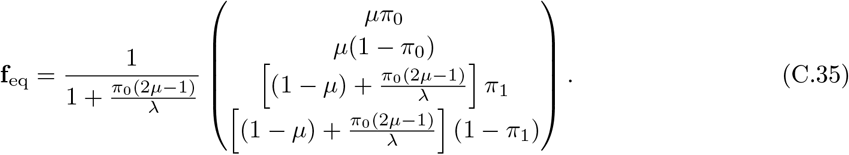

#### C.3 Per-genotype effective fitness and phenotypic buoying

The counterintuitive “surprise” that the genotype 1, phenotype 1 (red node in Main Text) beats genotype 0, phenotype 0 (blue node) despite having lower mapping probability and fitness is actually well-justified by ProP Gen theory. Recall that classical selection dynamics with phenotype noise during replication in the ProP Gen equations can be condensed into a collective term “PrGP selection”:

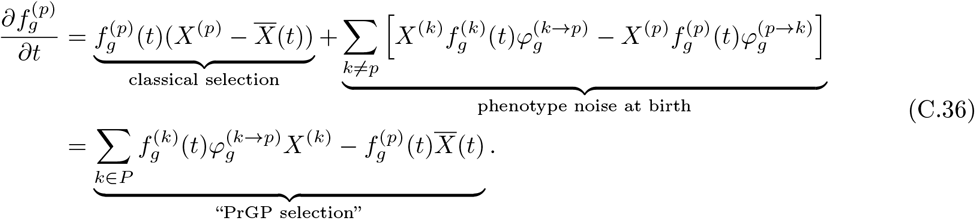

Summing over all phenotypes *p*, we can write a differential equation for the genotype frequencies:

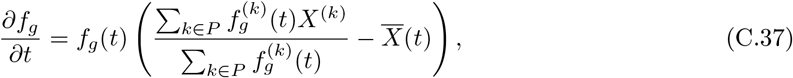

which simply looks like classical selection with a frequency and time-dependent per-genotype “effective fitness” given by the ratio within the parentheses above

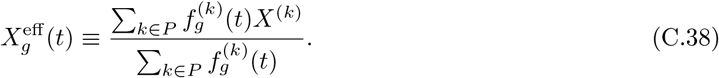

This shows that, for each genotype, PrGP selection makes a genotype-only fitness landscape appear *time-dependent*, and the genotype’s growth rate is exactly the per-genotype average fitness. Thus, a positive feedback loop ensues where genotype 1, phenotype 0 (green node), which is high fitness and high probability, helps genotype 1 overall absorb more population density, which of course accelerates the acquisition of more population density. As time goes by, the effective fitness of genotype 1 increases rapidly, and the red node benefits from the green node shunting its population density to the red node via phenotype noise.

#### C.4 Exact phase diagrams for equilibrium frequency distributions for phenotypic buoys with 2 genotypes and 2 phenotypes

In the DGP classical limit, suppose there are two alleles. In the infinite population limit (no genetic drift) with no mutations, where only selection dominates the dynamics, the allele with higher fitness will approach frequency 1 as the system approaches equilibrium, while the allele with lower fitness will approach frequency 0. With mutations, it is possible for both alleles to coexist at equilibrium. With PrGP maps, we showed analytically that, in general, multiple phenotypes can coexist at equilibrium, even when there are no mutations. We calculated the equilibrium frequencies of GP pairs in the two genotype, two phenotype scenario in eq. (C.23).

Here, we exactly calculate theoretical phase diagrams of coexistence between the four possible GP pairs in the two genotype, two phenotype scenario. The “phases” are relative orderings (permutations) of the equilibrium frequencies of the four GP pairs, assuming that no two frequencies are perfectly equal. There are 4! = 24 possible orderings. To more conveniently explore the phase diagram in a two-dimensional visualization, we select the probability of genotype 1 mapping to phenotype 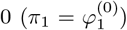 and the mutation rate (*µ*) to be the controllable variables in the phase diagram. We fix the phenotype fitnesess *X*^(0)^ = 0.09 and *X*^(1)^ = 0.02. We note that, since 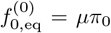 and 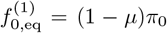, fixing *π*_0_ guarantees an equilibrium ordering of 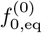 and 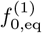. We will also fix *π*_0_ to three different values (*π*_0_ = 0.4 *<* 0.5, *π*_0_ = 0.5, and *π*_0_ = 0.6 *>* 0.5) in order to explore the shape of the (*π*_1_, *µ*) phase diagram in different *π*_0_ regimes. This reduces the maximum possible number of phases visible on the (*π*_1_, *µ*) phase diagram to 4!*/*2 = 12, though the real number may be less than 12 based on other constraints.

The phase boundaries can be computed by finding the conditions when any two of the equilibrium frequencies are equal. There are 4 frequencies, so that means there are 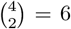 equations to consider. We handle each case below, ignoring the normalization constant common to all terms:

1. 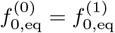, which is

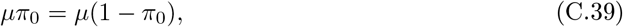

which simplifies to

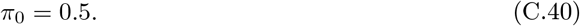

This curve will thus not be found on the (*π*_1_, *µ*), but we can expect that the shape of the (*π*_1_, *µ*) may change depending on the value of *π*_0_, which we will illustrate soon.
2. 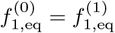, which is

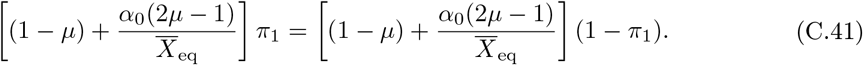

which simplifies to

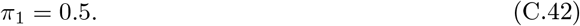

This curve will appear as a vertical line in the (*π*_1_, *µ*) phase diagram.
3. 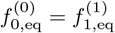, which is

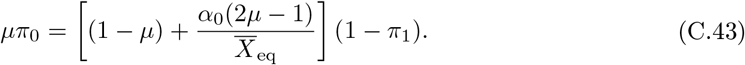

This will be an implicitly defined curve in the (*π*_1_, *µ*) space which can be solved numerically for plotting.
4. 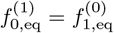, which is

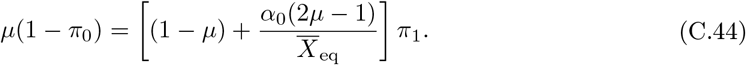

This will be an implicitly defined curve in the (*π*_1_, *µ*) space which can be solved numerically for plotting.
5. 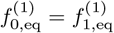, which is

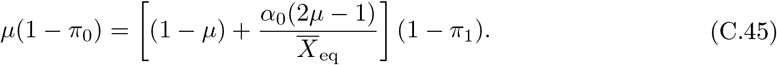

This will be an implicitly defined curve in the (*π*_1_, *µ*) space which can be solved numerically for plotting.
6. 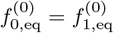, which is

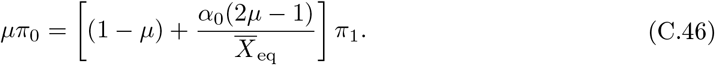

This will be an implicitly defined curve in the (*π*_1_, *µ*) space which can be solved numerically for plotting.

**Figure S1:**
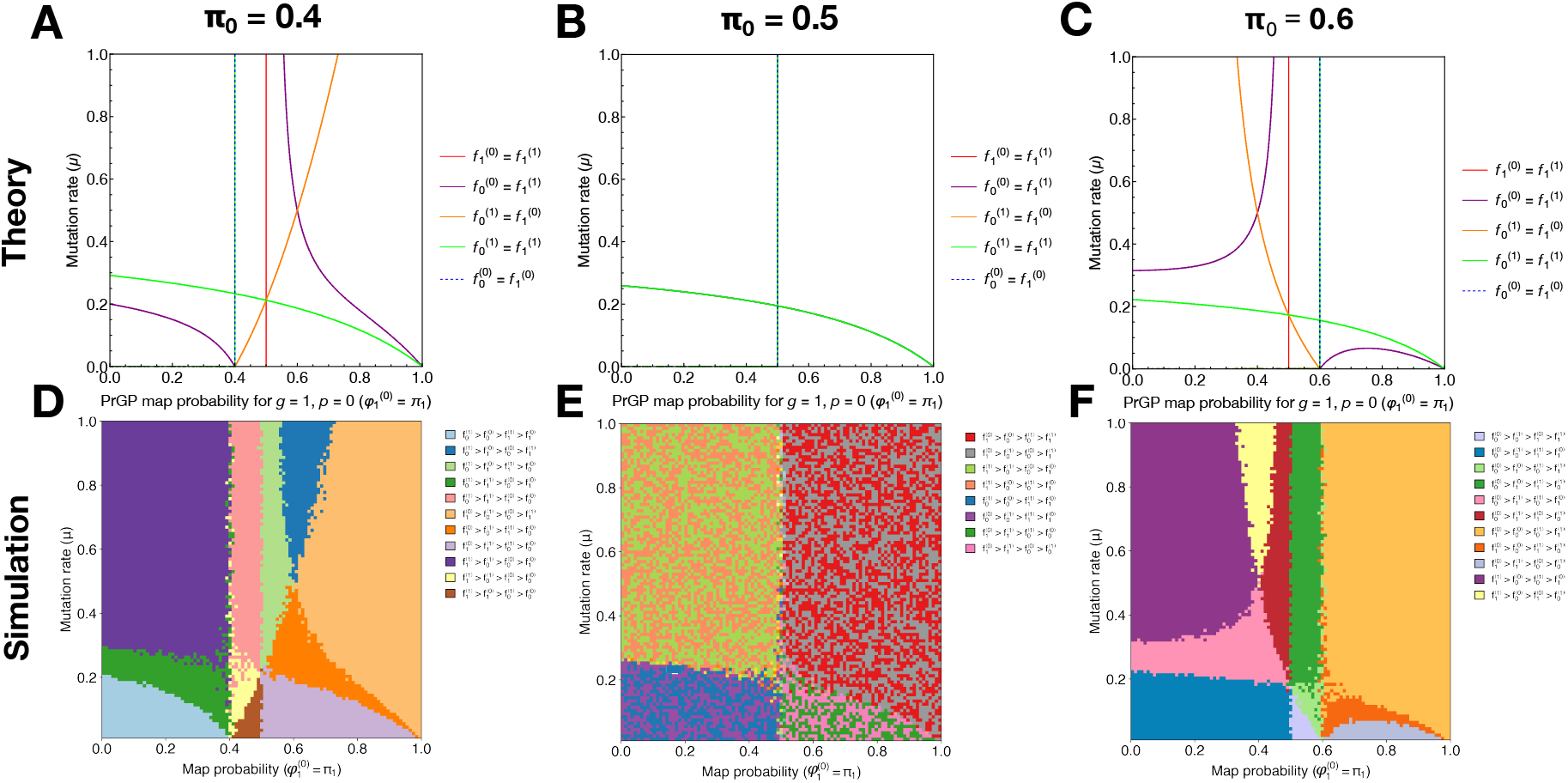
Complex phase diagrams of coexistence between genotype-phenotype pairs at equilibrium. (**A**-Theoretical phase boundaries for genotype-phenotype pair orderings at equilibrium for *π*_0_ = {0.4, 0.5, 0.6}. (**D**-**F**) Corresponding Numerical phase diagrams from `ProSeD` for *π*_0_ = {0.4, 0.5, 0.6}.

### D Phenotypic bridges

We now develop the theory of *phenotypic bridges*. Fitness valleys in the presence of DGP mapping require either operating in sufficiently polymorphic regime such that the population can spread out over multiple genotypes and eventually discover a distant peak, or it requires stochastic tunneling. Thus, valley crossing can be a very slow process especially when mutation rates are low or when the valley is very deep. To analytically investigate valley crossing in the presence of a “phenotypic bridge,” we consider three genotypes: 0, 1, and 2, and two phenotypes: 0 and 1. We assume that genotypes 0 and 2 deterministically map onto phenotype 0 (i.e. with probability 1), and that genotype 1 either maps onto phenotype 0 with probability *π* or onto phenotype 1 with probability 1 − *π*. We also assume that phenotype mapping probabilities do not depend on the initial phenotype, so 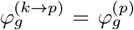. Thus, we have 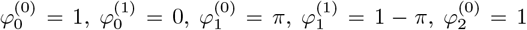, and 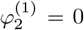. We also define shorthand labels for each of the 4 relevant genotype-phenotype pairs: the starting node has frequency 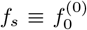, the bridge node has frequency 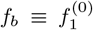, the valley node has frequency 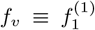, and the ending node has frequency 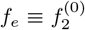. The mutation rates are determined by first selecting offspring to mutate with probability *µ*, and then a mutational neighbor is chosen with uniform probability, leading to a mutation matrix elements *µ*_0→1_ = *µ*_2→1_ = *µ* and *µ*_1→0_ = *µ*_1→2_ = *µ/*2. We assign fitnesses *X*^(0)^ = *X*_0_ and *X*^(1)^ = *γX*_0_, with 0 ≤ *γ <* 1. Now, we work in the diffusion limit with infinite population limit, assuming mutation probability at birth to be *µ*, and additionally only considering phenotype noise at birth:

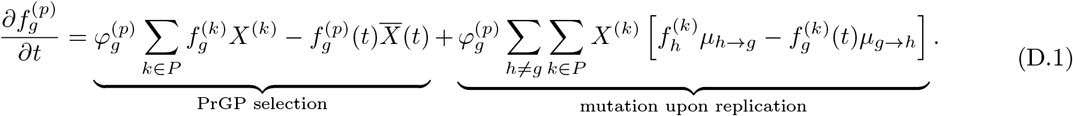

For each of the four genotype-phenotype pairs, we can write the differential equations

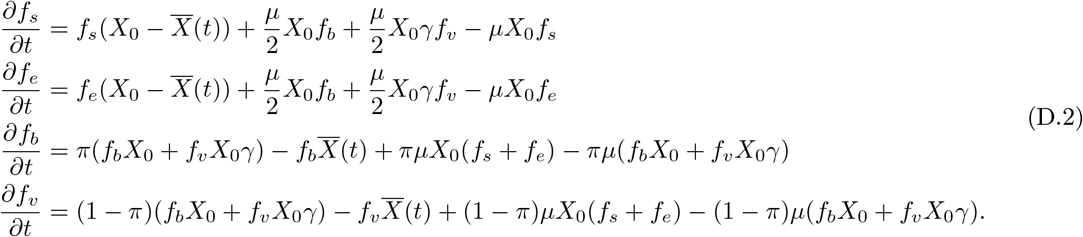

#### D.1 Exact equilibrium distribution

At equilibrium, the time derivatives vanish, and we can take the mean fitness-dependent terms to the left-hand side of the equation, setting up an eigenvalue equation for the equilibrium frequency vectors:

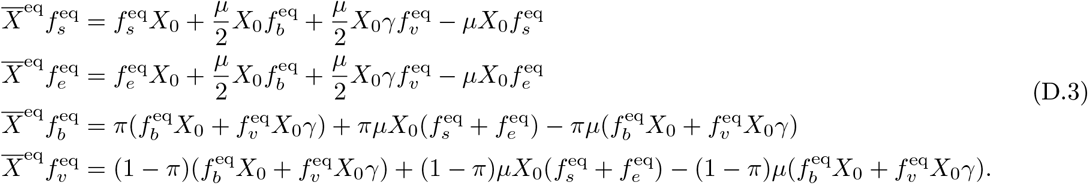

Diving the third and fourth equations, we immediately find

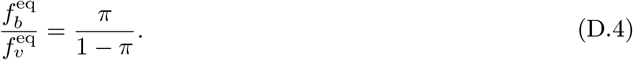

From the first two equations (and by symmetry of the setup), we can also see

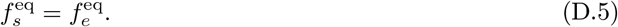

We also note that all equations can be divided by *X*_0_, to yield equations independent of *X*_0_, since

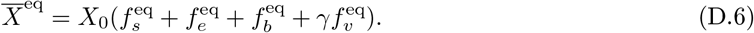

Defining

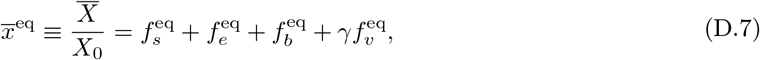

and

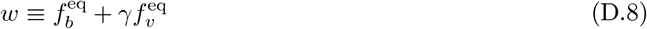

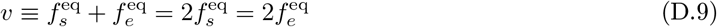

and making additional rearrangements, we now have

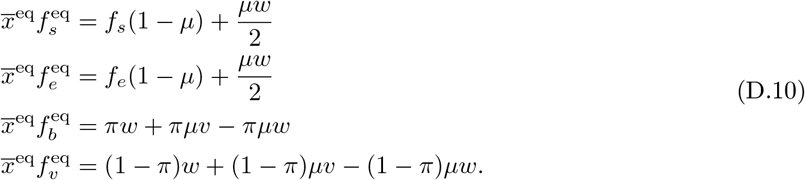

Adding the first two equations, we have

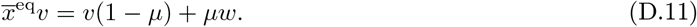

Then adding the third equation to the fourth equation multiplied by *γ*, we have

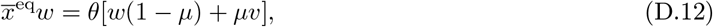

where we have defined

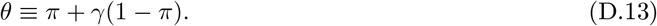

We can now write eq. (D.11) and eq. (D.12) as a 2 × 2 matrix equation

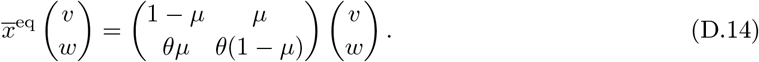

The characteristic equation is

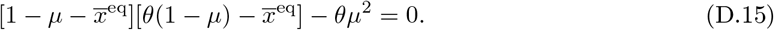

Solving the quadratic equation for 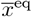 and taking the positive solution, we have

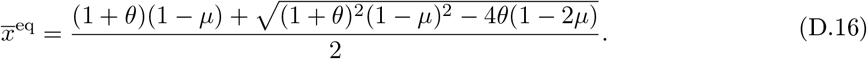

We can now rearrange eq. (D.11) to write

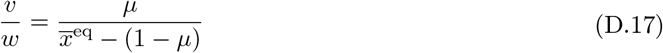

From eq. (D.4), we can write

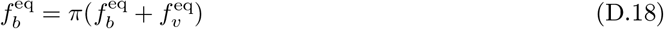

and

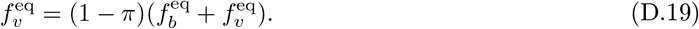

Now, it follows that

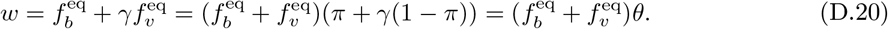

Thus, we can write

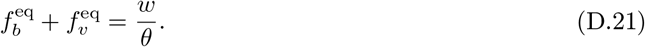

Now, we can use normalization of all four genotype-phenotype pair frequencies to write

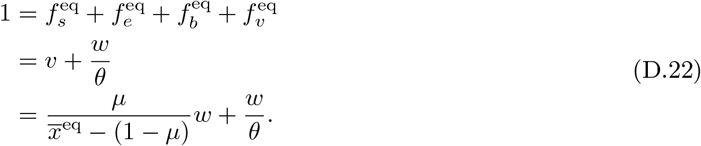

Solving for *w*, we have

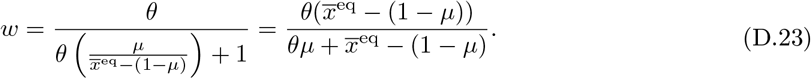

We can also write *v*:

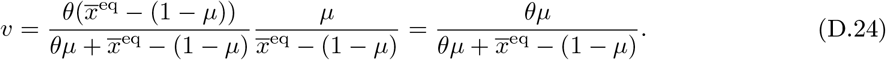

We can now write the equilibrium frequency vector, using 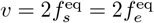 and 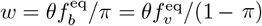:

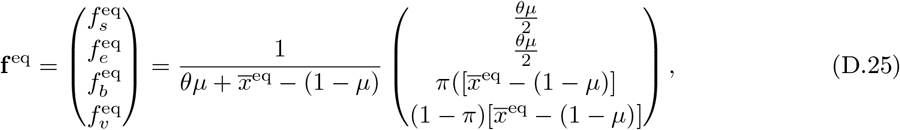

where 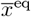 is given by eq. (D.16).

#### D.2 Relaxation time constant

We now linearize the nonlinear diffusion equations in eq. (D.2) around the equilibrium frequency vector and compute the time constant for the slowest decay mode toward equilibrium. Expanding the nonlinear equation around the equilibrium vector **f** = **f** ^eq^, we obtain

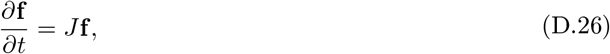

where *J* is the Jacobian

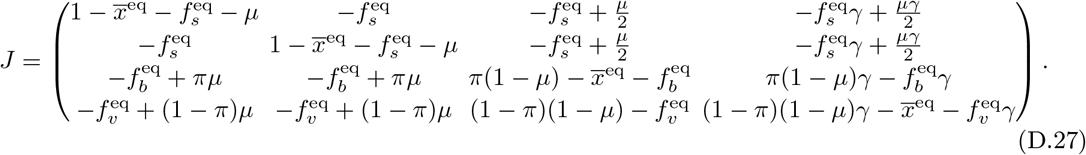

Due to normalization, one of the eigenvalues will be 0. Numerical evidence from the main text makes it clear that the bridge and valley genotype-phenotype pairs rapidly equilibriate while the slowest decay mode is the relaxation of *f*_*s*_ and *f*_*e*_, which appear to “mirror” each other while decaying toward exponential—that is, *f*_*s*_ and *f*_*e*_ experience equal but opposite changes while relaxing toward equilibrium. By inspection, it is readily verifiable from the Jacobian above that the vector

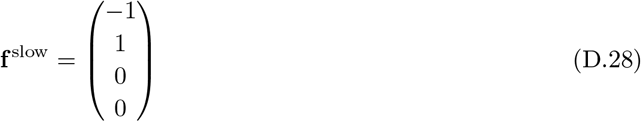

is indeed an eigenvector of the Jacobian. The corresponding eigenvalue,

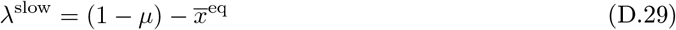

provides the exponential decay rate of the slowest mode. The associated exponential time constant for equilibrating over the bridge is

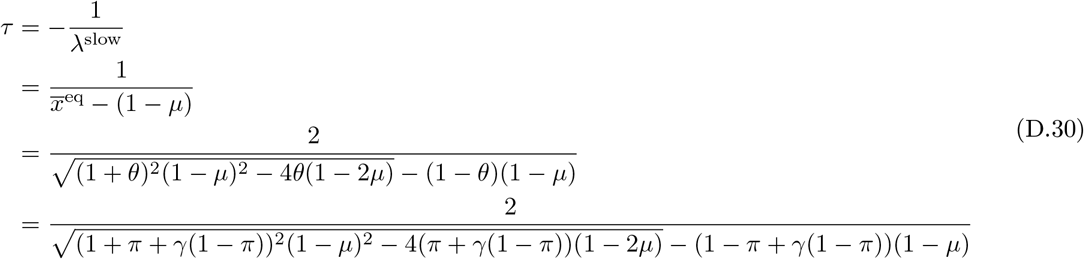

where we used *θ* = *π* + *γ*(1 − *π*), as defined earlier. We show in the main text that this theoretical time constant displays excellent agreement with numerical fits from exponentially fitting trajectory data.

### E Selection dynamics: global translational symmetry breaking of the fitness landscape

For the remainder of this supplementary material, we assume that the phenotype probability does not depend on the starting phenotype, so 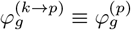, but rather only depends on the destination phenotype.

#### E.1 Global translational symmetry breaking of the fitness landscape

We now consider the selection term in the DGP and PrGP cases and show that the uncertain GP mapping in the latter case leads to dependence of the evolutionary dynamics on absolute, not simply, relative fitnesses. This also occurs when comparing the Eigen quasispecies equations with Kimura’s diffusion limit with mutations; the former couples mutations to fitnesses, while the latter treats mutations as an additive generator. Here, phenotype noise during replication is coupled to fitness while stochastic phenotype switching is an additive contribution.

The DGP selection term (i.e. the replicator equation) is:

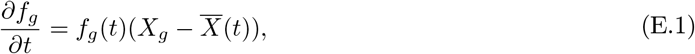

and the PrGP selection term (which we showed previously is the same as the DGP selection term plus a fitness-dependent phenotype switching term) has dependence on the GP mapping probability 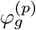:

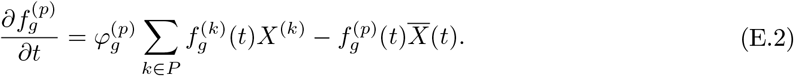

We now consider a shift of the entire fitness landscape by some absolute fitness *A*, so that all fitnesses are translated additively:

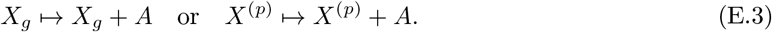

The DGP selection term becomes:

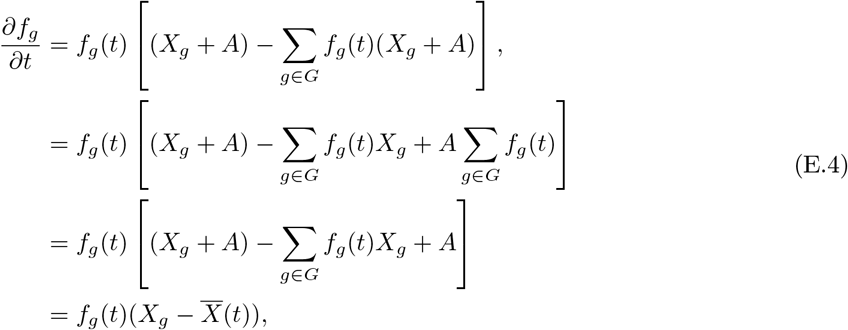

which means it does not transform at all. Akin to how only relative differences in potential energy impact trajectories in classical mechanics, only relative differences in growth rates affect evolutionary dynamics in the deterministic GP mapping case, as is well known. The PrGP selection term, however, demonstrates a major difference:

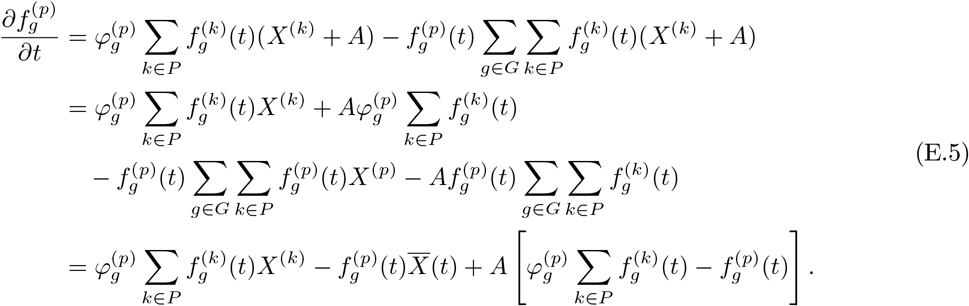

Thus, in the PrGP case there remains a dependence on *A*, indicating that *absolute* fitness, and not simply relative fitness (as is the case in canonical population genetics), determines evolutionary dynamics.

#### E.2 Mean fitness can decrease with time

Moreover, we now construct an example to show that it is possible for the change in mean fitness to be *negative*. We are searching for a class of solutions where

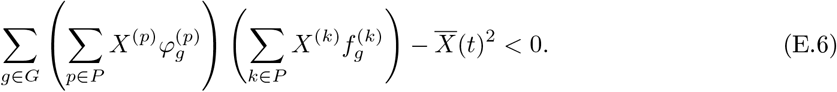

Consider a system with two genotypes and two phenotypes, each marked with indices 0 and 1. Suppose we have fitnesses

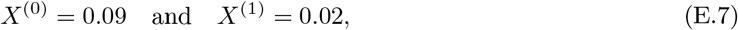

GP mapping probabilities

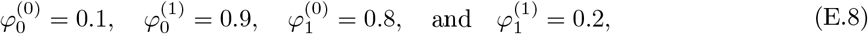

and initial frequencies

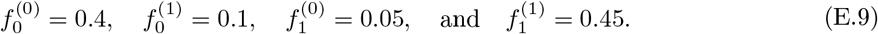

The mean fitness is

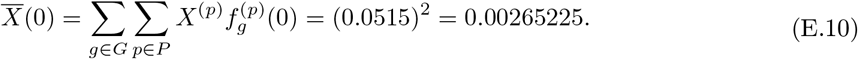

The first term in the expression eq. (E.6) is

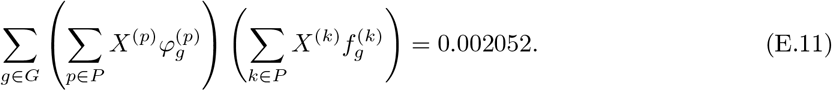

Since 0.002052 *<* 0.005776, we have successfully found an example where

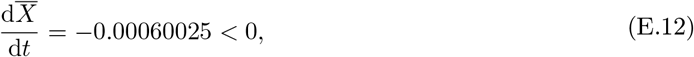

which means that mean fitness can decrease *only with selection* dynamics. We simulate this particular case with the initial conditions given above in Figure S2C-E. A summary of the probabilities, fitnesses, and genotype network topology is shown in Figure S2C. Frequency trajectories can show non-monotonicity (Figure S2D). Genotype 0, phenotype 1 (shown in orange), has high GP mapping probability but is low fitness. Thus, asymptotically, its frequency will be low. But, initially, its frequency grows because of the relatively high initial frequency of genotype 0, phenotype 0 (blue). The rapid growth of the genotype 0, phenotype 1 and non-monotonic behavior contributes to the initial decrease in mean fitness before its eventual increase towards its equilibrium value (Figure S2E).

Over the last 90 years, many have argued against the validity Fisher’s fundamental theorem of natural selection, suggesting that misinterpretation of the theorem, mutations, environmental interactions, epistasis, linkage disequilibrium, and other factors can prevent mean fitness from increasing with time. We have now shown that probabilistic mapping of genotype to phenotype can also cause decrease in mean fitness. Unlike classical DGP selection, where genotype frequencies tend towards fixation if they have the uniquely highest fitness, in PrGP selection, high fitness phenotypes can still “feed” lower fitness phenotypes. We discuss this phenomenon, which we call “phenotype buoys,” in depth in the following section.

### F Exact persister cell resuscitation dynamics

Here, we construct ProP Gen theory equations for phenotype noise and SPS associated with phenotypic resuscitation dynamics. We then show that they are exactly solvable. In the experiments of Fang and Allison^36^, they allow persister cells to resuscitate after washing away antibiotics. The dormant persister cells (P) can then resuscitate into three active phenotypes: failed (F), damaged (D), and healthy (H). Dormant persisters only resuscitate but they do not replicate until they have transitioned to an active phenotype, so we set the dormant fitness to zero *X*^(*P* )^. The damaged and failed phenotypes both are morphologically aberrant, but the failed ones cannot replicate, so we set their fitness to zero *X*^(*F* )^ = 0. Damaged active cells can still replicate with fitness *X*^(*D*)^ which is presumed to be less than or equal to the healthy cells’ fitness *X*^(*H*)^.

The transition rate (which is an SPS rate *S*(*t*)) out of the dormant persister state is empirically measured to be exponential in time^36^, which is highly suggestive of a positive feedback mechanism. They empirically validate the form:

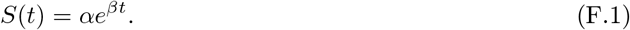

**Figure S2:**
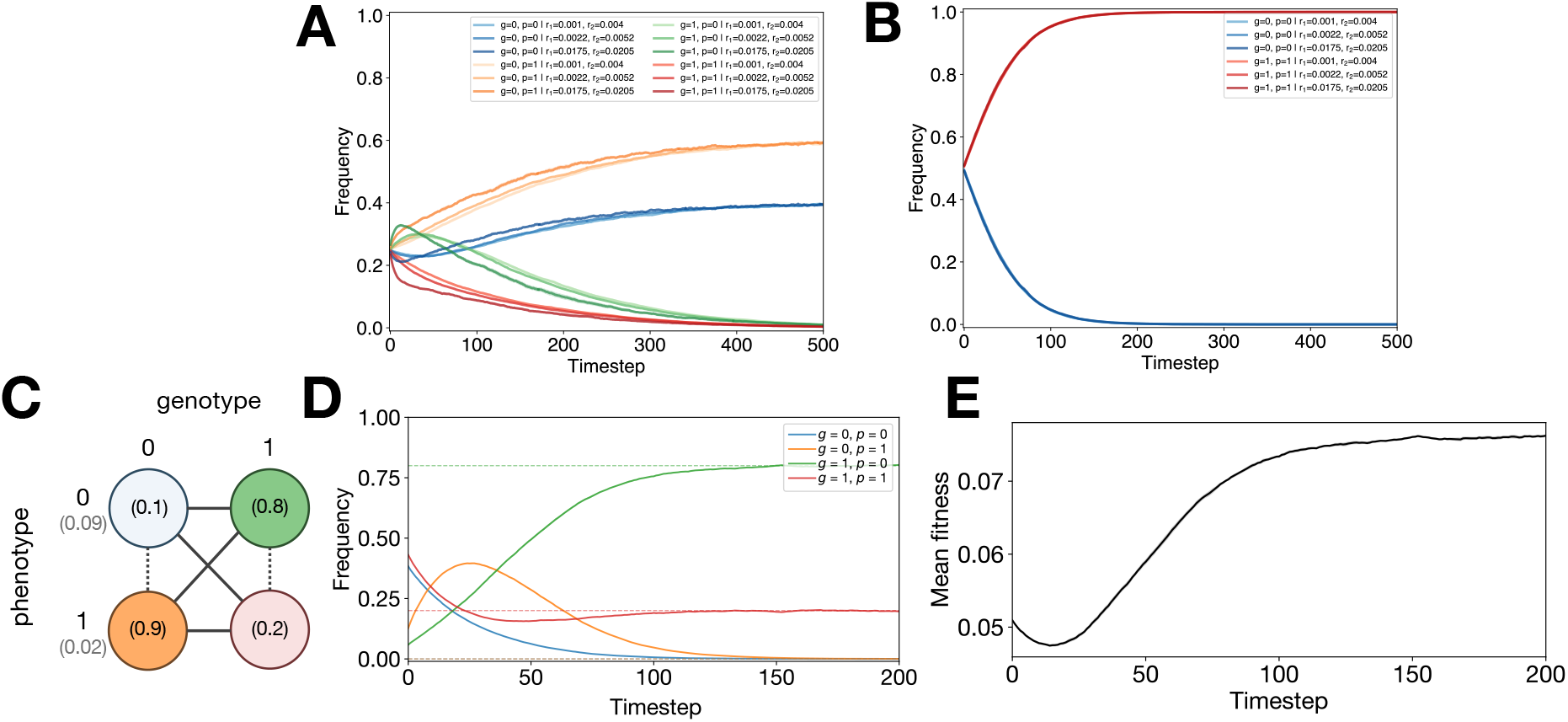
Phenotypic uncertainty creates a dependence on absolute fitness. (**A**) PrGP dynamics translated by different absolute fitnesses while keeping relative fitnesses the same. Translational symmetry breaking is observed. (**B**) DGP dynamics at different absolute fitnesses. Translational invariance is observed. (**C**) Diagram of two genotype, two phenotype set up for the PrGP cases. (**D**) `ProSeD` simulation of the system with the specified initial frequencies from Section C.2. (**E**) Mean fitness versus time.

Thus, the number of persister cells decays as an exponential of an exponential. We call the SPS probabilities out of the dormant state *σ*^(*P* →*i*)^ for *i* ∈ {*D, F, H*}, and ∑_*i*_ *σ*^(*P* →*i*)^ = 1. Furthermore, the damaged state’s cell division (“partitioning”) can be with compromised fidelity, either creating more damaged cells or creating healthy cells. This is a key example of phenotype noise at birth, in contrast to SPS. Thus, we have *φ*^*D*→*H*^ as the probability that the offspring of a damaged cell is healthy, and *φ*^*D*→*D*^ = 1 − *φ*^*D*→*H*^ is the probability that the offspring of a damaged cell is damaged.

Given the phenotypic transitions described above, we can now write down the ProP Gen equations for the four phenotypes. However, it is also important to note that Fang and Allison^36^ did not impose any population bottleneck (i.e. via serial dilution), so if we were to redo the derivation from Section A, we would not have a mean fitness term imposing selection due to Malthusian fitnesses. We also will not include the effect of sampling noise via genetic drift for the same reason. Further fluctuations would be due to noise in the parameters themselves such as fitnesses and the mapping or SPS probabilities. Thus, we write down the ODEs for absolute abundances {*n*_*i*_(*t*)} the four phenotypes:

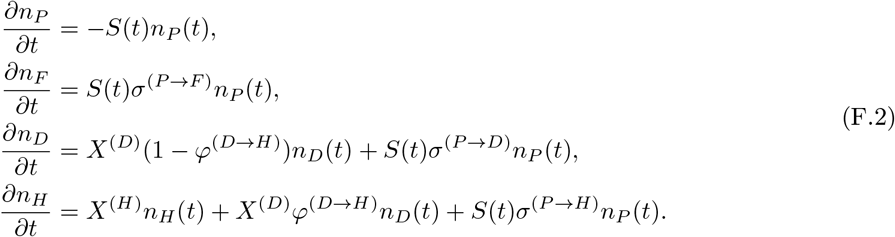

We now will show that these equations admit an exact solution in terms of special functions. Since the healthy cells depend on healthy, damaged, and persister cells; the damaged cells depend on damaged and persister cells; the failed cells depend on the failed cells; and the persister cells only depend on persister cells, it will make most sense to solve the equations in order for *P* → *F, D* → *H*.

First, we note the persister cell dynamics are solvable from separation of variables, as done by Fang and Allison^36^:

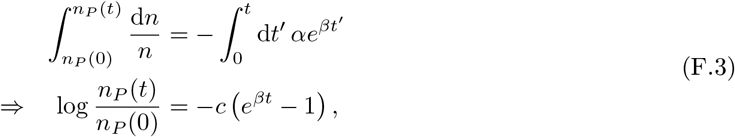

where we have defined *c* ≡ *α/β* for future notational convenience. Rearranging, we have

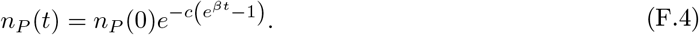

Now, substituting the first line of eq. (F.2), into the second line, we find

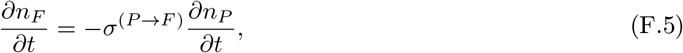

which we then integrate:

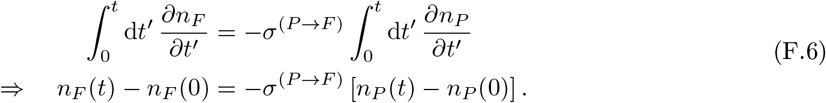

Rerranging, this gives us

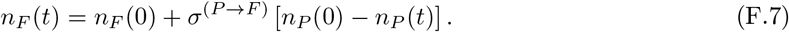

Next, we define *a* = *X*^(*D*)^ 1 − *φ*^(*D*→*H*)^ for convenience and rearrange the third line of eq. (F.2) to obtain

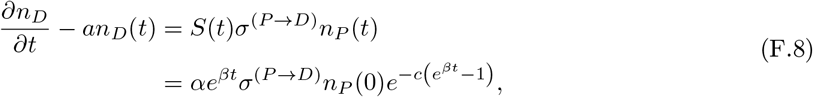

where in the second line we have substituted in the previously solved result for *n*_*P*_ (*t*). Now, we note that

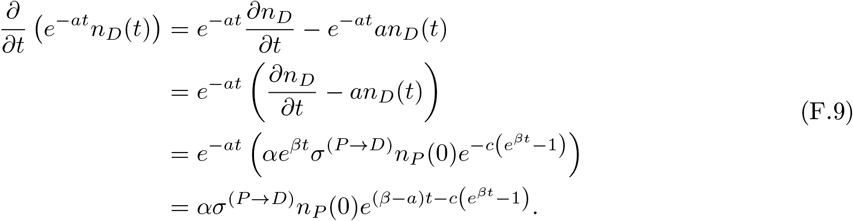

Integrating both sides, we have

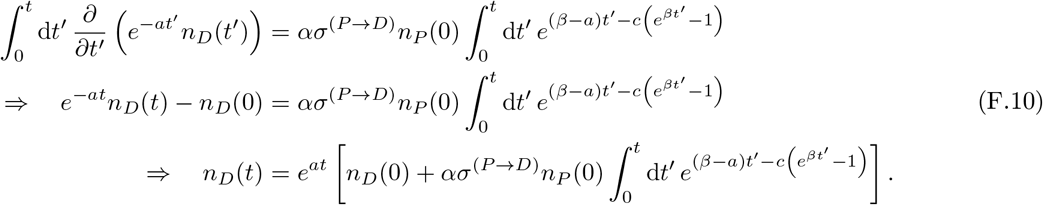

Now, we evaluate the integral. First, we perform the *u* substitution by setting 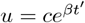, which gives us d*u* = *βu* d*t*^*′*^ or d*t*^*′*^ = d*u /*(*βu*). We also have *e*^*βu*^ = *u/c* and *t*^*′*^ = *β*^−1^ log(*u/c*). It then follows that

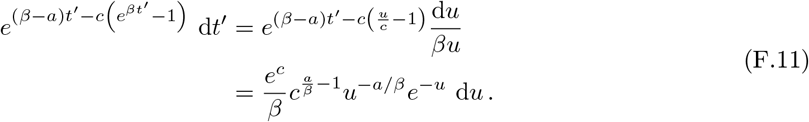

Now, we can rewrite the integral

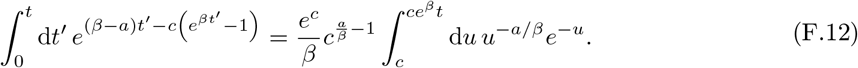

If we let *q*_*D*_ = 1 − *α/β*, we can rewrite the integral as

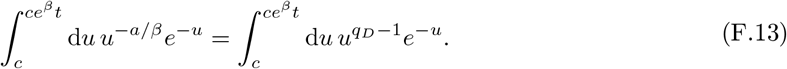

Using the definition of the lower incomplete gamma function

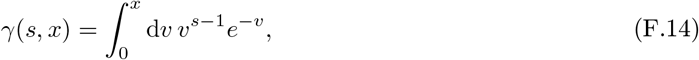

we can now write

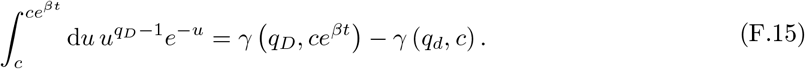

Thus, we have

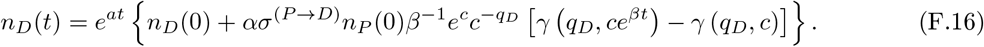

Finally, we turn to the dynamics of the healthy cells. We define a linear combination of the healthy and damaged cells

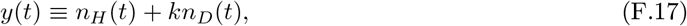

which, upon differentiation, gives

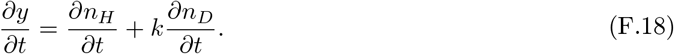

Substituting in the equations from eq. (F.2), this becomes

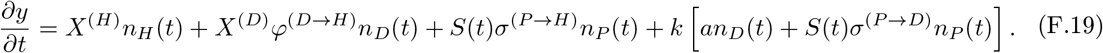

Substituting back in *n*_*H*_ (*t*) = *y*(*t*) − *kn*_*D*_(*t*) and simplifying, we have

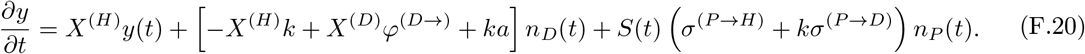

If we can make the bracket vanish, then 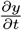 only depends on terms which can be written again in terms of the lower incomplete gamma function. So, we try to find the *k* which makes the bracket vanish

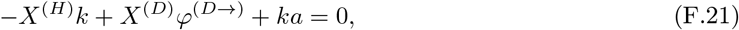

which yields

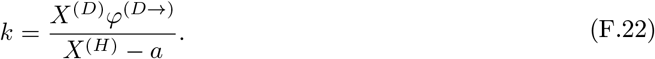

Defining *B* ≡ *σ*^(*P* →*H*)^ + *kσ*^(*P* →*D*)^ for convenience, we can now write

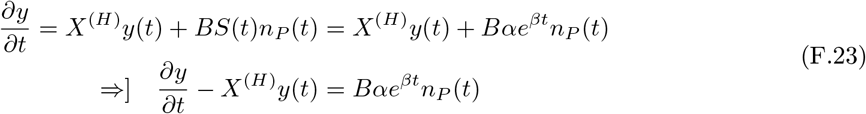

Now, noting that

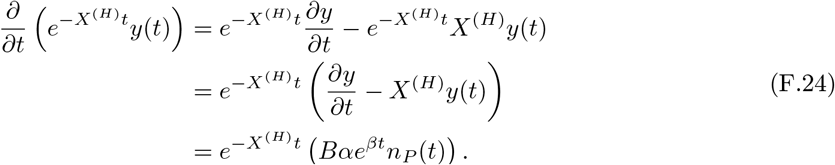

Integrating both sides, we have

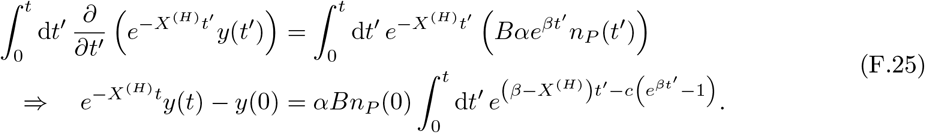

We can solve the integral using the same *u* substitution as before, except now we have *X*^(*H*)^ in place of *a*. So, defining *q*_*H*_ ≡ 1 − *X*^(*H*)^*β*, we can write the integral as

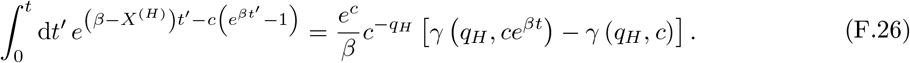

Finally, we have

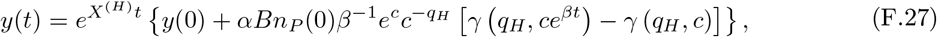

with *y*(0) = *n*_*H*_ (0) + *kn*_*D*_(0). Now, replacing *y*(*t*) using the solutions to *n*_*D*_(*t*) and eq. (F.17), we have the abundance of healthy cells

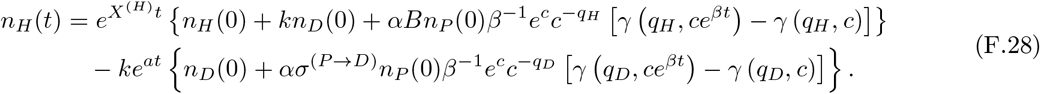

